# Regnase-1 Promotes Tumor-Initiating Activity in Non-Small Cell Lung Cancer

**DOI:** 10.1101/2025.07.08.663650

**Authors:** Keito Okazaki, Madoka Kawaguchi, Shohei Murakami, Haruna Takeda, Hiroki Sekine, Osamu Takeuchi, Hozumi Motohashi

## Abstract

Regnase-1, encoded by the *ZC3H12A* gene, is a well-known RNase that suppresses inflammation by degrading the mRNAs of inflammatory cytokines. However, its role in cancer pathogenesis, especially in non-small cell lung cancer (NSCLC), remains poorly understood. Through an analysis of public databases, we found that NSCLC patients with higher *ZC3H12A* expression levels had a worse prognosis than those with lower levels. To explore the function of Regnase-1 in NSCLC, we knocked out the *ZC3H12A* gene in NSCLC cell lines and compared their transcriptomes with those of parental cells. This analysis identified the SOX2 pathway as a common pathway suppressed by Regnase-1 deficiency. Consistent with the SOX2 contribution to the cancer stemness, Regnase-1 inhibition impaired oncosphere growth and tumor formation of cell lines derived from adenocarcinoma, squamous cell carcinoma and large cell carcinoma. It was also effective for NRF2-activated NSCLC cells, which are highly resistant to most of the therapeutics. Notably, post-tumorigenic suppression of Regnase-1 significantly inhibited tumor growth, suggesting that Regnase-1 could be a promising therapeutic target for post-tumorigenic treatment of NSCLC. Given recent studies describing that Regnase-1 inhibition enhances anti-cancer immunity, we propose that targeting Regnase-1 could be an ideal strategy for controlling intractable cancers by both suppressing cancer cells and activating anti-cancer immunity.

## Introduction

Non-small cell lung cancer (NSCLC) is the most common form of lung cancer, representing approximately 85% of all cases. Within NSCLC tumors, a small subpopulation of cells, known as cancer stem cells (CSCs), play a pivotal role in promoting disease progression, metastasis, therapeutic resistance, and recurrence^1,2,3^. The significance of cancer stemness in NSCLC cannot be overstated, as it directly contributes to the aggressive nature and poor prognosis of the disease. CSCs exhibit a high resistance to conventional treatments such as chemotherapy and radiation, often surviving these therapies and driving tumor relapse and further metastasis. Recent studies suggest that CSCs may play a role in the failure of immune checkpoint inhibitor therapy^4^. Targeting molecular drivers in CSCs, alongside standard treatments, has the potential to enhance patient outcomes and achieve long-lasting therapeutic responses. CSCs exhibit multiple characteristics, some of which directly confer resistance against conventional therapies. These include enhanced drug detoxification and efflux, cell cycle quiescence, reliance on fatty acid oxidation in mitochondria, and the expression of unique cell surface antigens^5^. Based on these characteristics, various strategies have been developed to target CSCs, such as inhibiting drug efflux transporters, inducing cell cycle entry to promote differentiation, blocking lipolysis and mitochondrial function, and utilizing chimeric antigen receptor (CAR) T cell therapy to exploit CSC-specific antigens. In the context of NSCLC, antigens and signaling pathways unique to CSCs have been identified^6,7^, providing diagnostic biomarkers for the early detection of relapse and therapeutic targets for achieving complete disease remission. Efforts have been made to identify and explore therapeutic targets to combat CSCs in NSCLC, either by directly inhibiting their activity^8,9^ or by inducing their cell cycle entry and differentiation into conventional tumor cells that are more responsive to standard treatments^10^. Among the detrimental characteristics of CSCs, tumor-initiating activity is one of the critical contributors to tumor relapse and subsequent metastasis.

Regnase-1, which is encoded by the gene *ZC3H12A*, is well-known as an RNase that suppresses inflammation by degrading inflammatory cytokine mRNAs^11,12^. Recent studies have demonstrated important roles of Regnase-1 not only in inflammation control but also in various biological processes and pathological contexts. For instance, Regnase-1 plays a regulatory role in the lineage selection of hematopoietic stem cells, dictating the fate of lymphoid and myeloid differentiation^13^. In the field of oncology, several papers have described both beneficial and detrimental roles of Regnase-1 in cancer cells. Regnase-1 was shown to suppress breast cancer progression by degrading anti-apoptotic genes in its RNase activity-dependent manner^14^. In clear cell renal cell carcinoma, Regnase-1 was reported to inhibit epithelial-mesenchymal-transition (EMT) by suppressing the Wnt/β-Catenin signaling pathway^15^. Tissue-specific disruption of *Zc3h12a* in mice accelerated carcinogenesis in the respective tissue: pancreatic cancer by pancreas-specific disruption of *Zc3h12a*^16^ and skin cancer by epidermis-specific disruption of *Zc3h12a*^17^. A recent study showed that an RNase-dead Regnase-1 activates c-Met signaling, which in turn regulates reprogramming transcription factors such as Sox2, c-Myc, Klf4 and Oct4^18^, thereby promoting cancer stem-like phenotypes^19^. Another study reported that Regnase-1 targets and degrades a specific group of miRNAs, the loss of which inhibits cell cycle^20^. Collectively, these findings suggest that Regnase-1 suppresses cancer stemness and cell proliferation through its RNase activity. Taken together, these reports describe a role of Regnase-1 as a negative regulator of cancer development. In contrast, Regnase-1 has been reported to promote cell proliferation, migration and angiogenesis, leading to malignant progression of glioma and poor patient prognosis^21^.

Whereas the role of Regnase-1 in cancer cells appears to vary depending on the cellular context, inhibition of Regnase-1 in immune cells has been reported to enhance anti-tumor immunity. Adoptive cell therapy is a recently developed modality of immunotherapy, whose efficacy is closely linked to the long-term persistence and effector function of T cells. Regnase-1 deficiency in CD8+ T cells^22^ or chimeric antigen receptor-T cells^23^ was found to increase their persistence, expansion and potency, demonstrating that targeting Regnase-1 is a promising strategy to improve the efficacy of adoptive cell therapy. Deletion of Regnase-1 in NK cells has also been reported to enhance IFN-γ-mediated anti-tumor immunity^24^.

In our previous study, we searched for potential therapeutic targets of NRF2-activated NSCLC^9,25^. NRF2-activated cancers refer to NSCLC and other malignancies in which NRF2 is persistently and excessively activated due to a failure in its degradation mechanism. Because persistently activated NRF2 confers resistance to chemotherapy, radiotherapy and immunotherapy, NRF2-activated cancers are among the most challenging to treat^26,27^.

Regnase-1 was identified as one of the NRF2 target genes specifically in *KEAP1*-mutant NRF2-activated NSCLC cell lines^9^. Because Regnase-1 function in NSCLC had never been reported, and because Regnase-1-high NSCLC cases had a worse prognosis than Regnase-1-low cases in the TCGA database, we decided to investigate whether Regnase-1 promotes the malignant progression of NSCLC, expecting that, if so, Regnase-1 inhibition would be an ideal bifunctional therapy to suppress cancer cells and activate anti-tumor immunity.

We found that *ZC3H12A* disruption in NSCLC cell lines decreased SOX2 pathway genes, implying that Regnase-1 supports cancer stem-like phenotype. Consistently, oncosphere growth *in vitro* and tumorigenesis in serial xenograft experiments were suppressed by Regnase-1 inhibition in multiple NSCLC cell lines irrespective of their NRF2 activation status, suggesting that Regnase-1 promotes tumor-initiating activity. Post-tumorigenic inhibition of Regnase-1 was also effective to antagonize the tumorigenesis. Of note, NSCLC cell lines derived from squamous cell carcinoma (LUSC) and large cell carcinoma (LCC), for which effective targeted therapies have not been developed^28^, were also effectively suppressed by Regnase-1 inhibition. These results suggest that Regnase-1 is a possible therapeutic target for LUSC and LCC as well as intractable NRF2-activated NSCLC.

## Results

### NSCLC cases with high expression of *ZC3H12A* have a worse prognosis

In our previous study exploring potential therapeutic targets for NRF2-activated NSCLCs, we identified *ZC3H12A* as a gene directly regulated by NRF2 specifically in NRF2-activated NSCLC cells. Although we expected that *ZC3H12A* expression is higher in NRF2-activated NSCLCs than NRF2-normal NSCLCs, a correlation between NRF2 activity and *ZC3H12A* expression was not clear in either LUAD or LUSC when we analyzed the TCGA public database (Supplementary Fig. S1A). Consistently, Regnase-1 protein levels varied independently of NRF2 protein levels (Supplementary Fig. S1B). These observations indicate that the transcriptional regulation by NRF2 does not necessarily imply high levels of *ZC3H12A* expression and suggest the presence of additional regulatory factors for *ZC3H12A* expression.

In contrast, when we examined whether *ZC3H12A* expression levels correlated with the prognosis of NSCLC patients, we found that both LUAD and LUSC cases with high *ZC3H12A* expression had a worse clinical outcome than those with low *ZC3H12A* expression (Fig. 1). These observations suggest that Regnase-1 contributes to the malignancy of NSCLC irrespective of NRF2 activation status.

**Figure 1.**
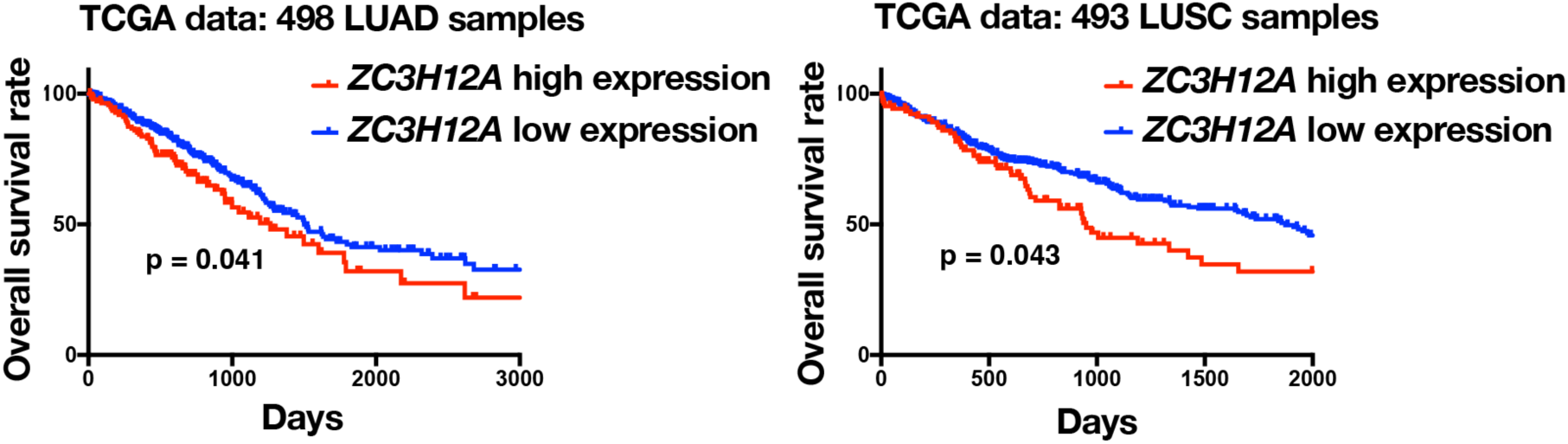
Correlation between *ZC3H12A* expression and patient outcomes in NSCLC. Overall survival rates for patients in the TCGA database grouped into *ZC3H12A* high expression cases and *ZC3H12A* low expression cases in lung adenocarcinoma (LUAD) and lung squamous cell carcinoma (LUSC). Kaplan-Meier analysis was performed. Statistical significance was evaluated by the log-rank test.

### Regnase-1 is associated with SOX2 and EMT pathway in NSCLC cell lines

To verify the function of Regnase-1, we disrupted *ZC3H12A* gene by CRISPR-Cas9 genome editing technology in two different NSCLC cell lines, A549 and H2023 (Supplementary Fig. S2). To exclude possible off-target effects, we established multiple mutant clones using two different guide RNAs (ΔZ cells) and confirmed reduction of Regnase-1 protein levels (Fig. 2A). Using each one of ΔZ cell clones established from A549 and H2023 cells and each parental cell, we performed RNA sequencing analysis to elucidate the Regnase-1-dependent transcriptome (Fig. 2B). Genes that were commonly downregulated or upregulated by *ZC3H12A* disruption in both cell lines were selected and subjected to Enrichr pathway analysis (Fig. 2C). SOX2 pathway, which is a transcription factor to maintain cell stemness – encompassing clonogenicity, pluripotency, and self-renewal^29^–was enriched in commonly downregulated genes, whereas EMT-related genes were enriched in commonly upregulated genes. Similar alterations in EMT-related gene expression were previously reported in clear cell renal cell carcinoma^15^.

**Figure 2.**
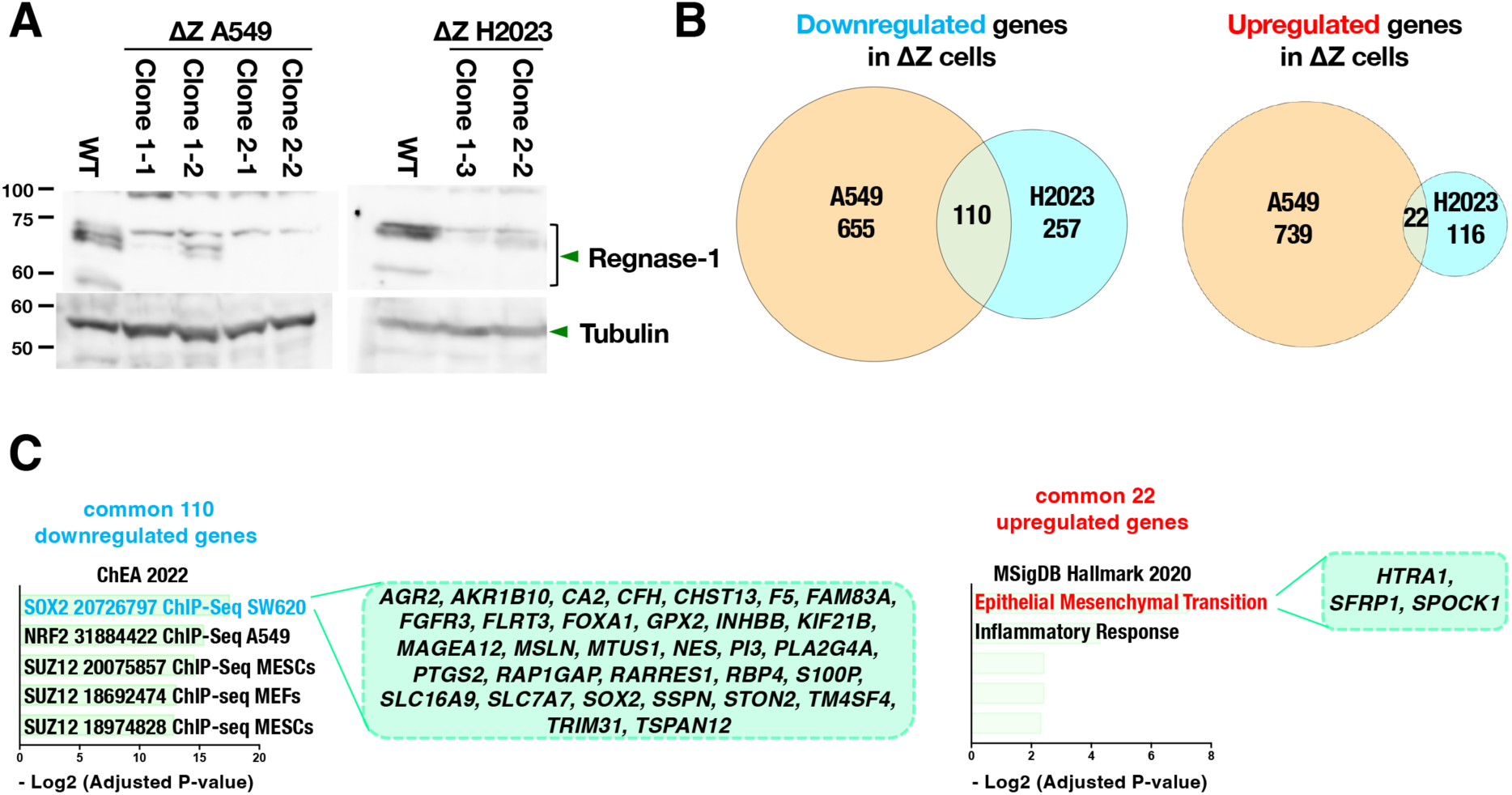
Analysis of Regnase-1-dependent transcriptome. (A) Immunoblot analysis detecting Regnase-1 in WT (wild-type) and *ZC3H12A*-deficient (ΔZ) cells established from A549 and H2023 cells. Tubulin was detected as a loading control. The result shown is a representative of 3 independent experiments. (B) Downregulated genes in ΔZ cells were defined as significantly downregulated genes (adjusted P value < 0.05) in Clone 2-2 ΔZ vs. WT of A549 cells and in Clone 2-2 ΔZ vs. WT of H2023 cells. 110 genes were identified as commonly downregulated genes between A549 and H2023 (left panel). Upregulated genes in ΔZ cells were defined as significantly upregulated genes (adjusted P value < 0.05) in Clone2-2 ΔZ vs. WT of A549 cells and in Clone2-2 ΔZ vs. WT of H2023 cells. 22 genes were identified as commonly upregulated genes between A549 and H2023 (right panel). (C) Annotation analysis of the commonly downregulated and upregulated genes was performed using “Enrichr.” Pathways shown in light blue are significantly decreased in ΔZ cells, while those in red are significantly elevated. Statistical significance was defined as an adjusted P-value of < 0.05.

As *ZC3H12A* disruption downregulated SOX2 pathway genes and upregulated EMT-related genes in two different NSCLC cell lines, we next investigated whether these gene expression alterations led to corresponding phenotypic alterations.

### Regnase-1 promotes oncosphere growth of NSCLC cell lines in vitro

Sox2 is recognized as a critical factor for maintaining the self-renewal capacity of stem-like population in NSCLC cancers^30,31^. Given that the loss of Regnase-1 resulted in decreased expression of SOX2 pathway genes (see Fig. 2C), we hypothesized that Regnase-1-deficient ΔZ cells would exhibit diminished stem-like phenotype. To test this hypothesis, we investigated whether the loss of Regnase-1 attenuates tumor-initiating activity, one of the features of stem-like phonotype, by examining oncosphere formation in vitro.

In addition to A549 and H2023 cells, we established ΔZ cells in H2172 cells, one of the LUAD cell lines (Supplementary Fig. S3). Cell growth in regular culture and spheroid formation, both reflecting simple cell growth ability, and oncosphere culture were compared between WT and ΔZ cells in all three cell lines. While cell growth in regular culture and spheroid formation did not show consistent trends (Fig. 3A and 3B), oncosphere growth was generally reduced in ΔZ cells of all three cell lines (Fig. 3C). The RNA-seq analysis comparing transcriptomes of WT and ΔZ oncospheres showed that SOX2 pathway genes were enriched in the commonly downregulated genes by Regnase-1 deficiency in A549 and H2023 cells (supplementary Fig. S4A and S4B), which is consistent with the impaired oncosphere growth of ΔZ cells.

**Figure 3.**
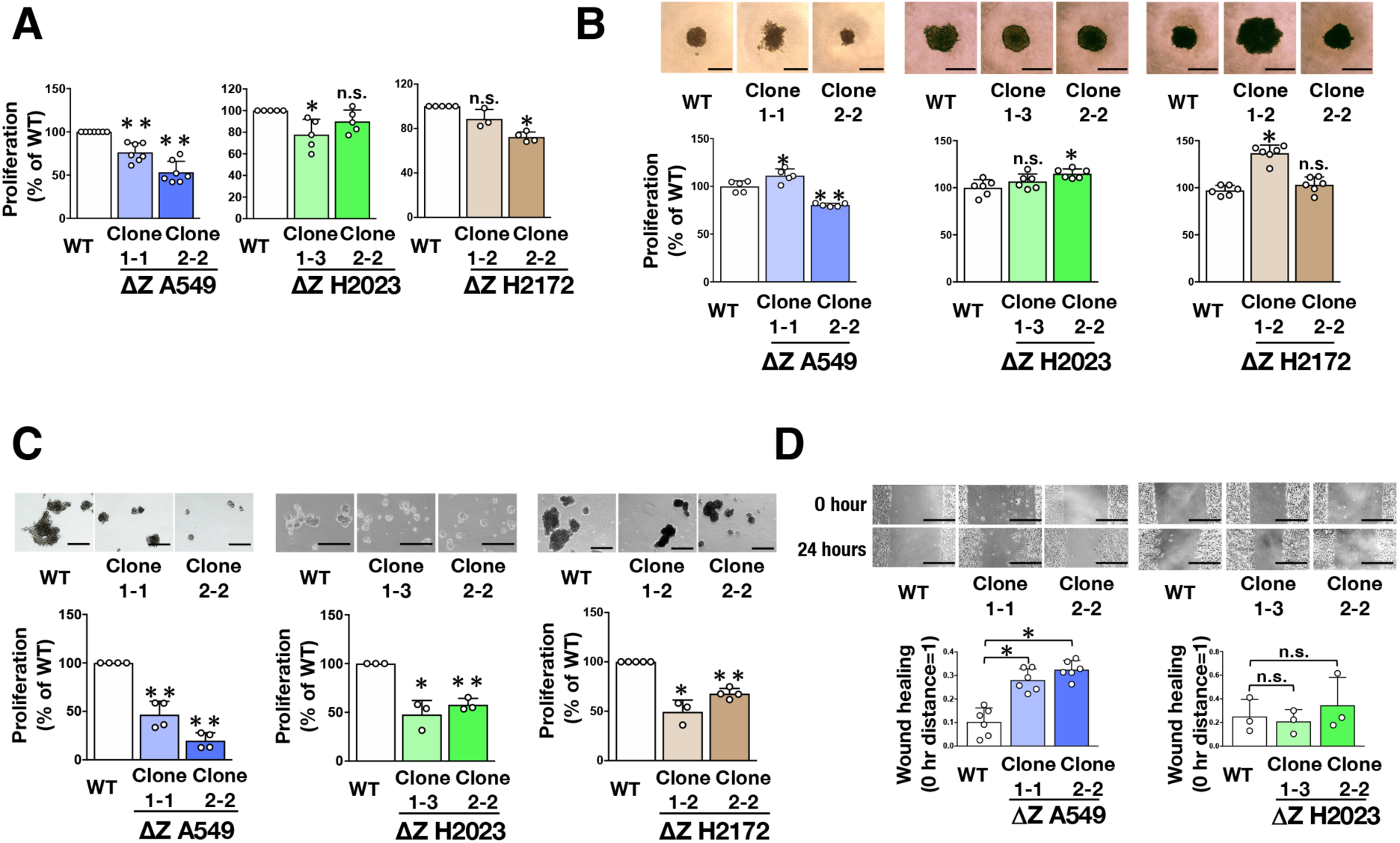
Regnase-1 promotes stem-like phenotype in NSCLC cell lines *in vitro*. (A) Cell proliferation of ΔZ cells generated from A549, H2023 and H2172 cells and their corresponding WT cells. A549, H2023 and H2172 cells (1 x 10^5^ cells) were seeded in low glucose DMEM supplemented with 10% fetal bovine serum on adherent culture dish. Cells were counted after 96 hrs for A549 cells, 72 hrs for H2023 cells, and 48 hrs for H2172 cells. Fold changes were calculated in comparison to WT. The average and SD of the fold changes from 3 independent experiments are shown. Two-sided confidence interval estimation was performed to evaluate statistical significance. * α<0.05, ** α<0.01, n.s.: not significant. (B) Day 4 spheroid growth of ΔZ cells generated from A549, H2023 and H2172 cells and their corresponding WT cells (upper panels). Scale bars indicate 100 μm. Cell numbers were estimated using a cell counting kit on day 4 (lower panels). Average cell numbers and SD from 5-6 independent experiments are shown. The average number of WT was set as 100% in each cell line. Two-sided Student’s *t* test was performed. \**p* < 0.05, \*\**p* < 0.005, n.s.: not significant. (C) Oncosphere growth of ΔZ cells generated from A549, H2023 and H2172 cells and their corresponding WT cells. Images of oncosphere growth (upper panels) and viable cell counts after trypsinization (lower panels). Fold changes were calculated in comparison to WT. Average cell numbers and SD from 3 or 4 independent experiments are shown. Two-sided Confidence interval estimation was conducted to evaluate statistical significance. * α<0.05, ** α<0.01. Scale bars indicate 500 μm. (D) Wound healing assay for evaluating motility of ΔZ cells generated from A549 and H2023 cells and their corresponding WT cells. Representative images of 0 and 24 hrs after scratching with a plastic pipette tip (upper panels) and quantification of wounded areas (lower panels). Average values and SD from 3 independent experiments are shown. Two-sided confidence interval estimation was conducted to evaluate statistical significance. * α<0.05, n.s.: not significant. Scale bars indicate 100 μm for A549 cells in (**C**) and 500 μm for the rest of the cells.

We also examined whether Regnase-1 inhibits EMT by performing a wound healing assay. Although a previous report described that reduction of Regnase-1 enhances EMT in clear cell renal cell carcinoma^15^, cell migration in the wound healing assay did not show consistent trends between A549 and H2023 (Fig. 3D). The ΔZ cells from A549 showed better cell migration than WT cells, but those from H2023 did not differ from WT cells, indicating that EMT inhibition by Regnase-1 is observed in limited cell lines derived from NSCLC.

We then evaluated in vivo tumor-forming ability of ΔZ cells. Tumorigenesis of ΔZ cells from all three cell lines was significantly suppressed in xenograft experiments compared to that of WT cells (Fig. 4A-4C), indicating that tumor-forming ability is impaired in ΔZ cells. Considering that ΔZ cells exhibited reduced oncosphere formation and diminished SOX2 pathway activity, we expected that their tumor-initiating activity would be compromised. A serial transplantation experiment using WT and ΔZ cells from A549 cells further suggested the impairment of the tumor-initiating activity by Regnase-1 deficiency (Fig. 4D and 4E). Together with the oncosphere growth results, we concluded that Regnase-1 promotes tumor-initiating activity in A549, H2023 and H2172 cells.

**Figure 4.**
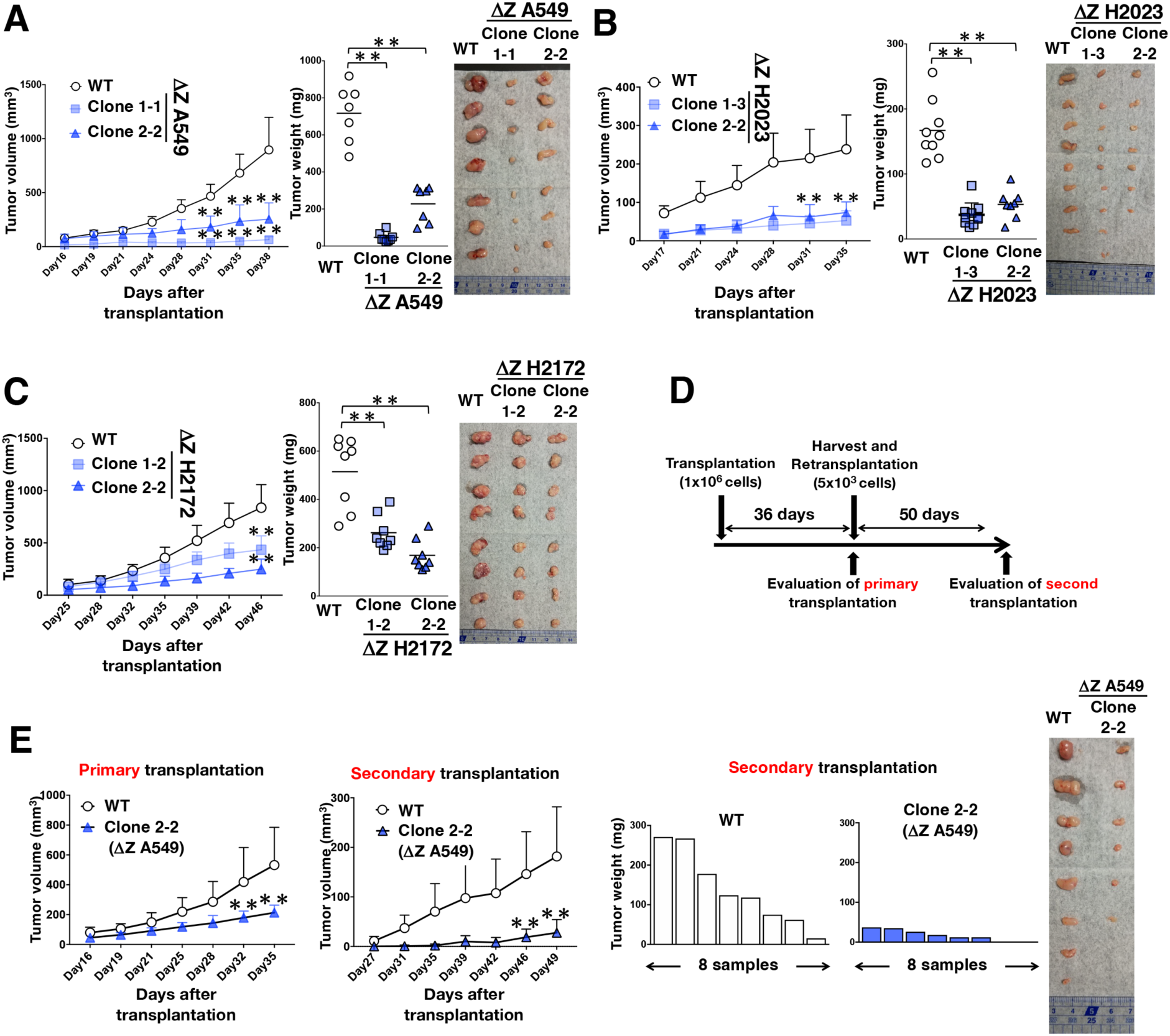
Regnase-1 promotes tumor-initiating activity in NSCLC cell lines *in vivo*. (**A-C**) Xenograft experiment of ΔZ cells generated from A549 (**A**), H2023 (**B**) and H2172 (**C**) cells and their corresponding WT cells (n=7-9 each; number of xenograft tumors). Photographs show xenograft tumors at the time of tumor weight measurement on day 38 (**A**), day 35 (**B**), day 46 (**C**). Data are presented as mean + SEM (left panels). Horizontal bars indicate the median tumor weight (middle panels). Two-sided Wilcoxon rank sum test was performed. \*\**p*<0.005. (**D, E**) Serial transplantation experiment of ΔZ and WT A549 cells. An experimental design (**D**) and growth of tumors (**E**). A photograph shows xenograft tumors at the time of tumor weight measurement in the secondary transplantation (**E**, right panel). In tumor growth curves, data are presented as mean + SEM (**E**, left two panels). Bar graphs show the weight of each tumor in the secondary transplantation of ΔZ and WT A549 cells (**E**, middle two panels). Two-sided Wilcoxon rank sum test was performed. \*\**p*<0.005.

### Regnase-1 promotes oncosphere growth in a broader spectrum of NSCLC cells

To evaluate the generality of Regnase-1-mediated promotion of tumor-initiating activity, we conducted transient *ZC3H12A* knockdown in 16 different NSCLC cell lines (Fig. 5A). Among them, 11 were derived from LUAD (Fig. 5B), 3 were from LUSC (Fig. 5C), and 2 were from LCC (Fig. 5D). All of these cell lines showed reduced oncosphere growth when *ZC3H12A* was transiently knocked down, except for 2 LUAD cell lines in which Regnase-1 reduction had no effect (Fig. 5B-5D). These results suggest that Regnase-1 contributes to the tumor-initiating activity in a broader spectrum of NSCLC cells. Of note, Regnase-1 inhibition was effective in reducing the oncosphere growth of NRF2-activated LUAD cell lines, except for H1944 (Fig. 5B, upper panels).

**Figure 5.**
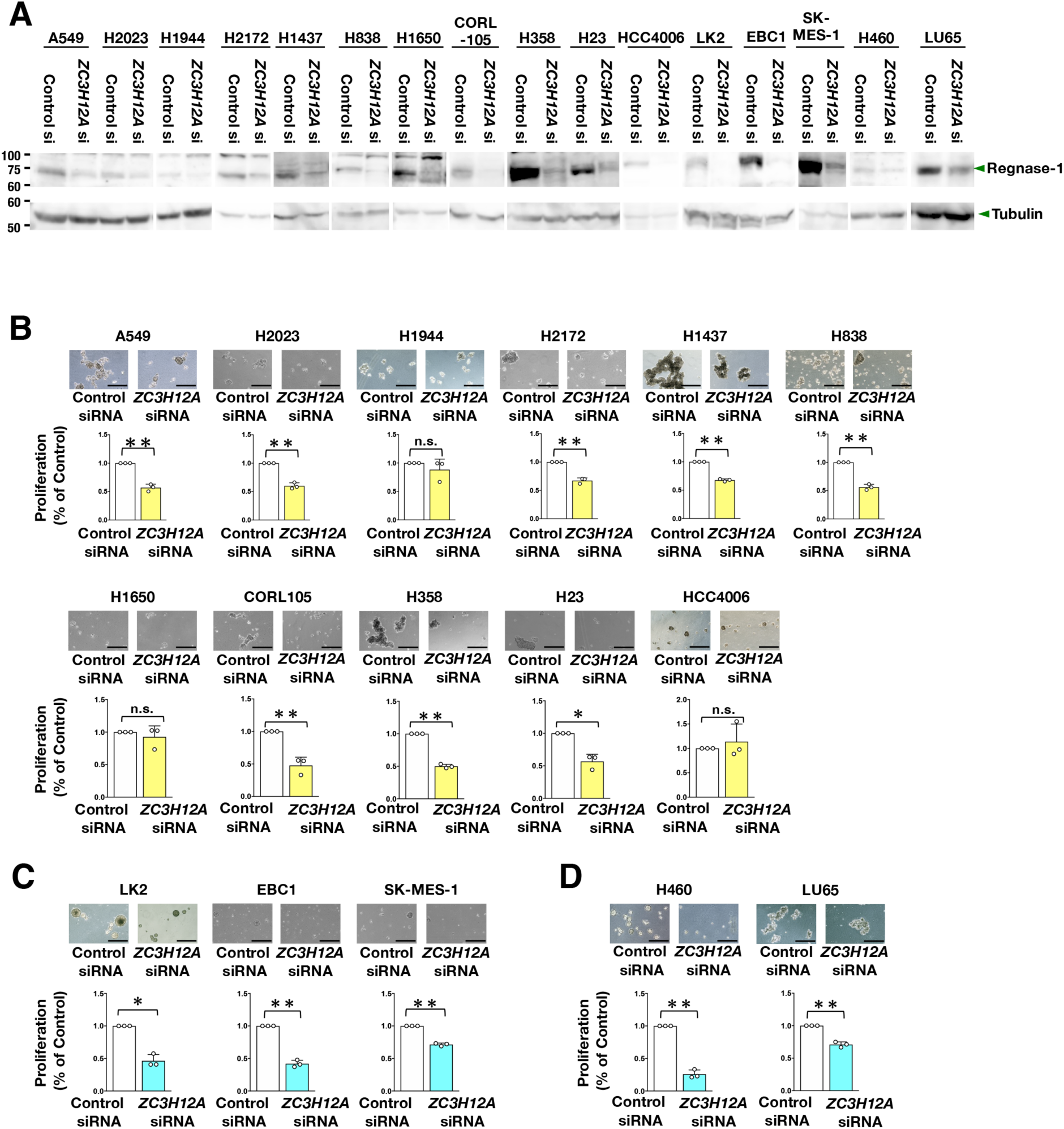
Regnase-1 contributes to stem-like phenotype in a broad range of NSCLC cells. (A) Immunoblot analysis detecting Regnase-1 levels in each NSCLC cell line treated with control siRNA and *ZC3H12A* siRNA. Tubulin was detected as a loading control. (**B-D**) Oncosphere growth of LUAD cell lines (**B**), LUSC cell lines (**C**) and large cell carcinoma cell lines (**D**) treated with control siRNA and *ZC3H12A* siRNA. Images of oncosphere growth treated with control siRNA and *ZC3H12A* siRNA (upper panels) and viable cell counts after trypsinization (lower panels). Fold changes were calculated in comparison to control siRNA. Average cell numbers and SD from 3 independent experiments are shown. Two-sided Confidence interval estimation was conducted to evaluate statistical significance. * α<0.05, ** α<0.01, n.s.: not significant. Scale bars indicate 100 μm for H1650 cells, 200 μm for H2172 cells and 500 μm for the rest of the cells.

A previous report described that overexpression of Regnase-1 in breast cancer cell line MDA-MB-231 induced apoptosis and suppressed tumor growth by degrading anti-apoptotic genes^14^. We were interested in an effect of Regnase-1 inhibition on tumor-initiating activity and conducted oncosphere culture of 3 breast cancer cell lines with and without *ZC3H12A* knockdown (Supplementary Fig. S5A). Although MDA-MB-231 oncosphere growth was not affected, the oncosphere growth of MCF7 and MDA-MB-468 was remarkably suppressed by *ZC3H12A* knockdown (Supplementary Fig. S5B). We also conducted transient *Zc3h12a* knockdown by two different siRNAs in murine melanoma cell line B16F10, which showed reduced oncosphere growth by *Zc3h12a* knockdown (Supplementary Fig. S5C and S5D). These data suggest a possibility that Regnase-1-mediated promotion of tumor-initiating activity is not restricted to NSCLC.

### Regnase-1 contributes to post-tumorigenic growth

Finally, we decided to investigate the efficacy of inhibiting Regnase-1 as a post-tumorigenic treatment. We used one of LCC cell lines H460 and one of LUSC cell lines LK2 for establishing cells with doxycycline (DOX)-inducible *ZC3H12A* knockdown (Fig. 6A). The effect of inducible *ZC3H12A* knockdown was first examined *in vitro* by oncosphere culture. DOX addition at the time of cell seeding (Fig. 6B) and at 4 days after the cell seeding (Fig. 6C) both reduced oncosphere formation. In xenograft experiments, *ZC3H12A* knockdown in H460 cells by DOX treatment before transplantation, followed by DOX treatment in the drinking water after transplantation, resulted in reduced tumorigenicity compared to the control (Fig. 6D). Importantly, *ZC3H12A* inducible knockdown in H460 cells after tumor formation effectively suppressed tumor growth compared to the control (Fig. 6E, upper right panels) but did not without DOX treatment (Fig. 6E, upper left panels). Similar results were observed in LK2 cells (Fig. 6E, lower panels). Inducible *ZC3H12A* knockdown in H460 cells exhibited comparable oncosphere growth and tumorigenic potential to control H460 cells without DOX treatment, allowing for a comparison between control and *ZC3H12A* knockdown cells with DOX treatment (Fig. 6E, upper panels). However, in LK2 cells, oncosphere growth and tumorigenic potential differed between control and *ZC3H12A* knockdown cells without DOX treatment, making it necessary to compare DOX-untreated and DOX-treated groups, respectively (Fig. 6E, lower panels). These findings highlight the potential of Regnase-1 as a promising therapeutic target.

**Figure 6.**
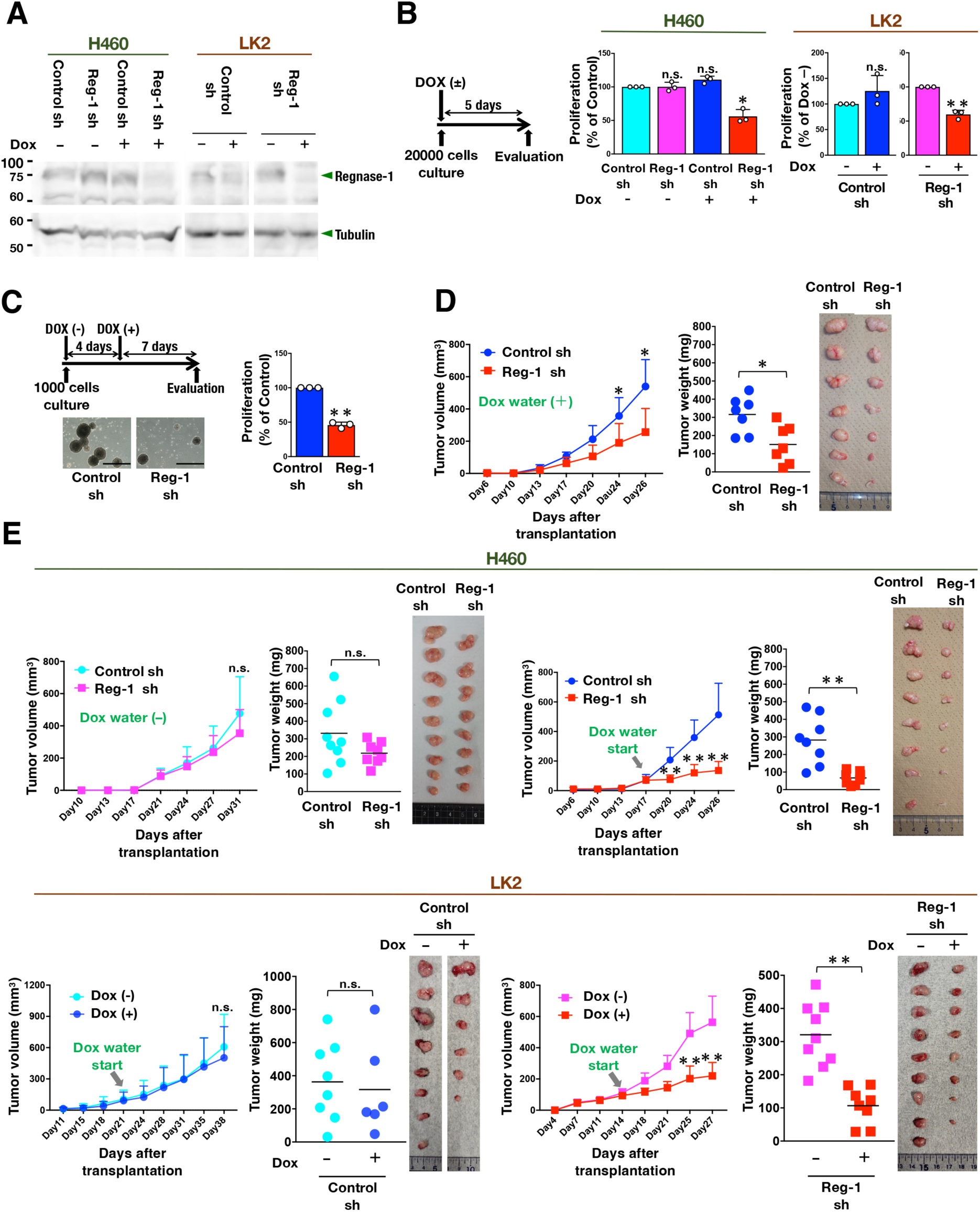
Regnase-1 contributes to post-tumorigenic growth (A) Immunoblot analysis detecting Regnase-1 levels in H460 and LK2 cells with inducible *ZC3H12A* knockdown (Reg-1 sh) and control H460 and LK2 cells (Control sh). The cells were treated with or without DOX (doxycycline) for 48 hours. Tubulin was used as a loading control. (B) Oncosphere growth of inducible *ZC3H12A* knockdown cells (Reg-1 sh) and control cells (Control sh) with Dox added at the start of cell seeding. Viable cells were counted after trypsinization. Fold changes were calculated in comparison to Control sh without DOX addition in H460 cells and Control sh or Reg-1 sh without DOX treatment in LK2 cells. Average cell numbers and SD from 3 independent experiments are shown. Two-sided Confidence interval estimation was conducted to evaluate statistical significance. * α<0.05, ** α<0.01, n.s.: not significant. (C) Oncosphere growth of inducible *ZC3H12A* knockdown (Reg-1 sh) and control (Control sh) in H460 cells with DOX added on day 4 after the cell seeding. Scale bars indicate 500 μm. Average cell numbers and SD from 3 independent experiments are shown. Two-sided Confidence interval estimation was conducted to evaluate statistical significance. ** α<0.01. (D) Xenograft experiments of H460 cells with inducible *ZC3H12A* knockdown (Reg-1 sh) and control H460 cells (Control sh) treated with DOX prior to 48 hrs of transplantation. The recipient mice were continuously fed with DOX water until the tumor evaluation on day 26. Photographs show xenograft tumors at the time of tumor weight measurement on day 26 (**D**). Data are presented as mean + SEM (left panels). Horizontal bars indicate the median tumor weight (middle panels). Two-sided Wilcoxon rank sum test was performed. \**p*<0.05. (E) Xenograft experiments of inducible *ZC3H12A* knockdown cells (Reg-1 sh) and control cells (Control sh) generated from H460 and LK2 cells. Each cell line was transplanted on day 0, and the recipient mice were fed with or without DOX water at the time indicated by the arrowhead. Photographs show xenograft tumors at the time of tumor weight measurement on day 31 or day 26 in H460 cells and day 38 or day 27 in LK2 cells. In the graphs showing tumor volume, data are presented as mean + SEM in the tumor volume. In the graphs showing tumor weight, horizontal bars indicate the median tumor weight. Two-sided Wilcoxon rank sum test was performed. \*\**p*<0.005, n.s.: not significant.

## Discussion

This study demonstrated that Regnase-1 enhances tumor-initiating activity across a broad spectrum of NSCLC cell lines, including those derived from LUAD, LUSC and LCC, and that its inhibition effectively suppresses post-tumorigenic growth in xenograft experiments. While many therapeutics exhibit varying efficacy across different subtypes of NSCLC, our results suggest that Regnase-1 may serve as a therapeutic target applicable to a wide range of NSCLC subtypes. Regarding two major subtypes of NSCLC, LUAD often exhibits actionable driving gene alterations, such as *EGFR* mutations and translocations involving *ROS1* and *ALK*, whereas LUSC rarely harbors actionable mutations^32^. LCC, which constitutes a small proportion of NSCLC, frequently demonstrates an aggressive tumor phenotype and lacks both targeted therapies and specific markers^33^. Consequently, LUSC and LCC pose significant treatment challenges due to their poor responsiveness to drugs and radiotherapy^34,35^, with no well-established molecular targeted therapies available for these subtypes. Even in LUAD, certain cases continue to experience suboptimal clinical outcomes despite significant advancements in molecular targeted therapies driven by cancer genome research^36^. This is often attributed to the presence of oncogenic mutations that complicate treatment efficacy^37^. A notable example is NRF2-activated LUAD, frequently associated with *KEAP1* mutations, often in combination with *KRAS* and *STK11* mutations^38^. These mutations confer extreme resistance to KRAS-targeted molecular therapies, even in the presence of a *KRAS* mutation^39^. NRF2-activated LUAD is also resistant to radiotherapy^40^ and immunotherapy^41^. We propose that targeting Regnase-1 represents a promising strategy to overcome these refractory forms of NSCLC.

From a cancer treatment perspective, Regnase-1 emerges as an ideal therapeutic target. Its inhibition in cancer cells suppresses tumorigenesis by reducing tumor-initiating activity, while in T cells, it enhances their ability to execute anti-cancer immunity^22–24^. We have previously reported that NOTCH3 inhibition suppresses tumorigenesis by reducing the tumor-initiating activity in NRF2-activated NSCLC^9^. Additionally, neutralizing antibodies against NOTCH3 have been shown to attenuate tumor growth of breast cancer cells^42^. However, blocking NOTCH3 also impacts vascular endothelial cells, leading to the promotion of tumor growth^43^. This suggests that while NOTCH3 antagonism may be effective in a cancer cell-autonomous context, it can be detrimental due to its effects on the tumor microenvironment. In contrast, inhibiting supersulfides, recently identified new entity of biomolecules, could be beneficial for both cancer cells and the tumor microenvironment, similar to the effects seen with Regnase-1 inhibition. Supersulfides, including persulfides and polysulfides, are upregulated under the control of NRF2^44^ and are frequently abundant in cancer cells, where they contribute to chemoresistance and exhibit anti-ferroptotic properties^45,46^. Additionally, supersulfides reduce inflammation by suppressing the proinflammatory M1 phenotype of macrophages^47,48^, which suggests that they may also inhibit anti-tumorigenic macrophage activity. Therefore, inhibiting supersulfides is expected to enhance the effectiveness of various therapeutic approaches, including chemotherapy, radiotherapy and immunotherapy. Therapies with such dual efficacy against both cancer cells and the tumor microenvironment represent the next generation of anti-cancer treatments, and Regnase-1 is highly likely to meet the criteria for such therapies.

In our *in vitro* experiments, there was no significant difference in the basic proliferation of WT and ΔZ cells (see Figures 3A and 3B). However, in *in vivo* experiments, the inhibition of Regnase-1 following tumor formation significantly suppressed tumor growth (see Figure 6E). These findings indicate that the ability of cells to proliferate *in vitro* is clearly distinct from their tumor-initiating activity, with Regnase-1 selectively contributing to the latter. Two major models of tumor development have been previously described, the clonal evolution model and the cancer stem cell model^49^. The former is a nonhierarchical model where mutations arising in tumor cells confer a selective growth advantage, whereas the latter is a hierarchical model where a small subset of cells has the ability to sustain tumorigenesis and generate heterogeneity through differentiation^49^. We consider that our results align with the latter. The anti-tumorigenic effects observed with the inhibition of Regnase-1 are of great clinical significance, suggesting that Regnase-1 may serve as a therapeutic target, not only for postoperative adjuvant chemotherapy but also in the treatment of advanced stages of NSCLC with anticancer drugs. Based on the phenotypes of Regnase-1-mutant mice^50–52^, potential side effects of Regnase-1 inhibition may include excessive immune activation, leading to autoimmunity similar to immune-related adverse events caused by immune checkpoint inhibitors. Therefore, once Regnase-1 inhibitors are developed, their dosage must be carefully optimized to prevent excessive immune activation.

An important unanswered question is whether the RNase activity of Regnase-1 contributes to its enhancement of cancer stem-like phenotype. To assess the necessity of RNase activity of Regnase-1, we attempted to rescue Regnase-1-deficient cells by reintroducing wild-type Regnase-1. However, despite using two different gRNAs, the Regnase-1-deficient clones showed only partial rescue in both oncosphere growth *in vitro* and tumorigenesis *in vivo* (data not shown). This partial rescue complicated the evaluation of whether an RNase-dead mutant, Regnase-1 D141N^53^, could fully restore the phenotypes of Regnase-1-deficient cells. During these experiments, we observed that both excessively low and high levels of Regnase-1 expression inhibited cell proliferation, and that regulating Regnase-1 protein abundance to endogenous levels was difficult. This difficulty in precise adjustment might have contributed to the failure in evaluating the activity of Regnase-1 D141N. A potential future experiment to assess the contribution of the Regnase-1 D141N mutation to the cancer stem-like phenotype could employ a doxycycline-inducible expression system, enabling precise control of Regnase-1 and its mutant expression levels. Alternatively, to ensure that the expression level of Regnase-1 D141N matches that of endogenous Regnase-1, we would either knock in the Regnase-1 D141N cDNA at the endogenous Regnase-1 (*ZC3H12A*) locus or use genome editing to replace Aspartate 141 with Asparagine in the endogenous Regnase-1 (*ZC3H12A*) locus. Aside from its RNase activity, some reports suggest that Regnase-1 may function as a transcription factor^54^. However, since it does not localize in the nucleus, this claim requires further verification. Nevertheless, if Regnase-1 operates independently of its RNase activity, it is plausible that it possesses an as-yet-undiscovered role, such as regulating intracellular signaling via protein-protein interactions, for instance.

Another remaining question is how Regnase-1 promotes the cancer stem-like phenotype. Contrary to a previous report^19^, the most probable effector downstream of Regnase-1 is SOX2. SOX2 is a crucial transcription factor with key roles in both normal and cancer stem cells. While controlling growth and proliferation of various normal stem cells^29^, SOX2 confers cancer stem cell properties, contributing to the self-renewal, maintenance of CSC populations, and tumor-initiating activity^30,31^. We found that Regnase-1-deficient ΔZ cells exhibited downregulation of SOX2 pathway genes and *SOX2* itself as well. We also observed partial recovery of *SOX2* expression and oncosphere growth when wild-type Regnase-1 was reintroduced into Regnase-1-deficient ΔZ cells (data not shown). However, transient knockdown of Regnase-1 did not change expression levels of *SOX2* (data not shown) but still inhibited oncosphere growth, implying that mechanisms beyond the SOX2 also contribute to Regnase-1-mediated enhancement of the cancer stem-like phenotype. The Wnt, Notch and Hedgehog pathways are often aberrantly activated in lung cancers and contribute to the maintenance of CSCs^7^. Transcriptomic analysis of Regnase-1-dependent genes in A549 and H2023 cells indicated that Regnase-1 suppresses the expression of sFRP1, a soluble inhibitor of Wnt signaling^55^. This implies that Regnase-1 may activate Wnt pathway by downregulating this inhibitory factor. However, transcriptome data from clinical tumor samples in the TCGA database revealed no significant correlation between Regnase-1 and SFRP1 or other genes involved in these CSC-supporting pathways. Moreover, a previous study of clear cell renal cell carcinoma described that Regnase-1 suppresses the Wnt/β-Catenin signaling pathway^15^. Thus, the contribution of Regnase-1 to tumor-initiating activity may depend on specific cellular contexts.

For the development of Reganse-1 inhibitors, addressing these unanswered questions will provide valuable insights. If the enhancement of tumor-initiating activity by Regnase-1 depends on its RNase activity, small molecules targeting its enzymatic function could be developed. Alternatively, if Regnase-1-interacting proteins play a critical role, disrupting these interactions with small molecules may serve as an effective strategy. A more straightforward approach is to reduce the Regnase-1 protein level, which could be achieved by inhibiting the transcription of *ZC3H12A* gene, nullifying Regnase-1 mRNA by antisense oligonucleotides or siRNA, or promoting Regnase-1 protein degradation using a PROTAC-based strategy.

In conclusion, although there are limitations as described above, this study showed that Regnase-1 is responsible for the tumor-initiating activity in a wide range of NSCLCs, including refractory forms. Regnase-1 will also be a good therapeutic target in the TME, thus the development of Regnase-1 inhibitors is expected in the future.

## Methods

### Mice

Four-week-old male BALB/cAJcl-nu/nu mice (CLEA Japan) were used in this study. All animals were housed in air-conditioned room at an ambient temperature of 20-26 °C, humidity of 30-70% and 12-hr dark/light cycle. They were housed in specific pathogen-free conditions. For the xenograft experiment, recipient mice were anesthetized via isoflurane inhalation. Once a sufficient depth of anesthesia was confirmed, cancer cells were transplanted subcutaneously. Mice were sacrificed on the specified days post-transplantation via cervical dislocation. All experimental procedures were performed in accordance with the regulations of the *Standards for Human Care and Use of Laboratory Animals of Tohoku University*, the *Guidelines for Proper Conduct of Animal Experiments* by the Ministry of Education, Culture, Sports, Science, and Technology of Japan and ARRIVE guidelines.

### Disruption of *ZC3H12A* to establish each ΔZ cells

Two guide RNAs (gRNAs) located in the 5th exon of the *ZC3H12A* gene were designed to disrupt all 3 variants. Lentiviral vectors expressing these gRNAs together with Cas9 mRNA were constructed by inserting annealed oligoDNAs (Supplementary Table S1) into lentiCRISPR v2, which was a gift from Feng Zhang (Addgene plasmid # 52961; http://n2t.net/addgene: 52961; RRID:Addgene_52961)^56^. A549, H2023 and H2172 cells were infected with lentiviral particles with 12.5 μg/ml polybrene. After incubation for 24 hrs, the cells were re-plated in 10 cm dishes and incubated in selection medium containing 2 μg/ml puromycin. Single clones were selected using cloning rings (TOHO). DNA was purified from each clone, and the modified regions were amplified using the primer sets listed in Supplementary Table S2. The PCR products were cloned and sequenced to verify disruption.

### Wound healing assay

Wound healing assay was based on previous reports^57^. Confluent A549 and H2023 cells in 60 mm culture dishes were scratched with a plastic pipette tip and cultured for 24 hrs. Photographs were taken with a DMi1 microscope (Leica). The area of lacking cells was measured using Image J, and migration index was calculated using the following formula: 1-(area after 24 hrs / area at the beginning). Additional materials and methods information is described in Supplementary Methods Doc. S1.

## Supporting information

Uncropped blots

## Acknowledgments

We thank the Biomedical Research Cores of the Tohoku University Graduate School of Medicine and Institute of Development, Aging and Cancer for providing technical support.

## Disclosure

### Ethics declarations

– Informed Consent: N/A
– Registry and the Registration No. of the study/trial: N/A
– Approval of the research protocol by an Institutional Reviewer Board: N/A
– Animal Studies: This study has been approved by the Animal Ethics Committee of Tohoku University under the number 2024MdA-007-03.

### Author contributions

K.O. conducted the experiments, analyzed the data and wrote early draft of the paper. M.K. and S.M. conducted the experiments and analyzed the data. H.T. conducted bioinformatic analysis. H.S. analyzed and interpreted the data. O.T. generated and provided critical materials and analyzed and interpreted the data. H.M. designed the study, conducted the experiments, supervised the research, analyzed the data and wrote early draft of the paper.

### Data availability

RNA-seq data were deposited at NCBI GEO (Accession No. GSE277438, https://www.ncbi.nlm.nih.gov/geo/query/acc.cgi?acc=GSE277438).

## References

1. Pandya, P., Al-Qasrawi, D. S., Klinge, S. & Justilien, V. Extracellular vesicles in non-small cell lung cancer stemness and clinical applications. Front Immunol 15, 1369356, doi:10.3389/fimmu.2024.1369356 (2024).

2. Min, H. Y. & Lee, H. Y. Mechanisms of resistance to chemotherapy in non-small cell lung cancer. Arch Pharm Res 44, 146–164, doi:10.1007/s12272-021-01312-y (2021).

3. Suresh, R., Ali, S., Ahmad, A., Philip, P. A. & Sarkar, F. H. The Role of Cancer Stem Cells in Recurrent and Drug-Resistant Lung Cancer. Adv Exp Med Biol 890, 57–74, doi:10.1007/978-3-319-24932-2_4 (2016).

4. Clara, J. A., Monge, C., Yang, Y. & Takebe, N. Targeting signalling pathways and the immune microenvironment of cancer stem cells - a clinical update. Nat Rev Clin Oncol 17, 204–232, doi:10.1038/s41571-019-0293-2 (2020).

5. Turdo, A., et al. Meeting the Challenge of Targeting Cancer Stem Cells. Front Cell Dev Biol 7, 16, doi:10.3389/fcell.2019.00016 (2019)

6. Okudela, K., et al. Expression of the potential cancer stem cell markers, CD133, CD44, ALDH1, and β-catenin, in primary lung adenocarcinoma--their prognostic significance. Pathol Int 62, 792–801, doi:10.1111/pin.12019 (2012).

7. García Campelo, M. R., Alonso Curbera, G., Aparicio Gallego, G., Grande Pulido, E. & Antón Aparicio, L. M. Stem cell and lung cancer development: blaming the Wnt, Hh and Notch signalling pathway. Clin Transl Oncol 13, 77–83, doi:10.1007/s12094-011-0622-0 (2011).

8. Zheng, Y., et al. A rare population of CD24(+)ITGB4(+)Notch(hi) cells drives tumor propagation in NSCLC and requires Notch3 for self-renewal. Cancer Cell 24, 59–74, doi:10.1016/j.ccr.2013.05.021 (2013).

9. Okazaki, K. et al. Enhancer remodeling promotes tumor-initiating activity in NRF2-activated non-small cell lung cancers. Nat Commun 11, 5911, doi:10.1038/s41467-020-19593-0 (2020).

10. Zito, G., et al. Retinoic Acid affects Lung Adenocarcinoma growth by inducing differentiation via GATA6 activation and EGFR and Wnt inhibition. Sci Rep 7, 4770, doi:10.1038/s41598-017-05047-z (2017).

11. Matsushita, K. et al. Zc3h12a is an RNase essential for controlling immune responses by regulating mRNA decay. Nature 458, 1185–1190, doi:10.1038/nature07924 (2009).

12. Mino, T. et al. Regnase-1 and Roquin Regulate a Common Element in Inflammatory mRNAs by Spatiotemporally Distinct Mechanisms. Cell 161, 1058–1073, doi:10.1016/j.cell.2015.04.029 (2015).

13. Uehata, T. et al. Regulation of lymphoid-myeloid lineage bias through regnase-1/3-mediated control of Nfkbiz. Blood 143, 243–257, doi:10.1182/blood.2023020903 (2024).

14. Lu, W. et al. MCPIP1 Selectively Destabilizes Transcripts Associated with an Antiapoptotic Gene Expression Program in Breast Cancer Cells That Can Elicit Complete Tumor Regression. Cancer Res 76, 1429–1440, doi:10.1158/0008-5472.CAN-15-1115 (2016).

15. Gorka, J. et al. MCPIP1 inhibits Wnt/β-catenin signaling pathway activity and modulates epithelial-mesenchymal transition during clear cell renal cell carcinoma progression by targeting miRNAs. Oncogene 40, 6720–6735, doi:10.1038/s41388-021-02062-3 (2021).

16. Okabe, J. et al. Regnase-1 downregulation promotes pancreatic cancer through myeloid-derived suppressor cell-mediated evasion of anticancer immunity. J Exp Clin Cancer Res 42, 262, doi:10.1186/s13046-023-02831-w (2023).

17. Morisaka, H., Takaishi, M., Akira, S. & Sano, S. Keratinocyte Regnase-1, a Downregulator of Skin Inflammation, Contributes to Protection against Tumor Promotion by Limiting Cyclooxygenase-2 Expression. J Invest Dermatol 143, 731–739, doi:10.1016/j.jid.2022.11.007 (2023).

18. Li, Y., et al. c-Met signaling induces a reprogramming network and supports the glioblastoma stem-like phenotype. Proc Natl Acad Sci U S A 108, 9951–9956, doi: 10.1073/pnas.1016912108 (2011).

19. Marona, P. et al. The endonuclease activity of MCPIP1 controls the neoplastic transformation of epithelial cells via the c-Met/CD44 axis. Cell Commun Signal 23, 28, doi: 10.1186/s12964-025-02029-x (2025).

20. Lichawska-Cieslar, A. et al. MCPIP1 modulates the miRNA‒mRNA landscape in keratinocyte carcinomas. J Exp Clin Cancer Res 43, 290, doi: 10.1186/s13046-024-03211-8 (2024).

21. Wang, R. et al. MCPIP1 promotes cell proliferation, migration and angiogenesis of glioma via VEGFA-mediated ERK pathway. Exp Cell Res 418, 113267, doi:10.1016/j.yexcr.2022.113267 (2022).

22. Wei, J. et al. Targeting REGNASE-1 programs long-lived effector T cells for cancer therapy. Nature 576, 471–476, doi:10.1038/s41586-019-1821-z<otherinfo> (2019)</otherinfo>.

23. Zheng, W., et al. Regnase-1 suppresses TCF-1+ precursor exhausted T-cell formation to limit CAR-T-cell responses against ALL. Blood 138, 122–135, doi:10.1182/blood.2020009309 (2021).

24. Sun, X. et al. Deletion of the mRNA endonuclease Regnase-1 promotes NK cell anti-tumor activity via OCT2-dependent transcription of Ifng. Immunity 57, 1360–1377.e1313, doi:10.1016/j.immuni.2024.05.006 (2024).

25. Okazaki, K. et al. CEBPB is required for NRF2-mediated drug resistance in NRF2-activated non-small cell lung cancer cells. J Biochem 171, 567–578, doi:10.1093/jb/mvac013 (2022).

26. Okazaki, K., Papagiannakopoulos, T. & Motohashi, H. Metabolic features of cancer cells in NRF2 addiction status. Biophys Rev 12, 435–441, doi:10.1007/s12551-020-00659-8 (2020).

27. Hayashi, M., Okazaki, K., Papgiannakopoulos, T. & Motohashi, H. The Complex Roles of Redox and Antioxidant Biology in Cancer. Cold Spring Harb Perspect Med 14, a041546, doi:10.1101/cshperspect.a041546 (2024).

28. Niu, Z., Jin, R., Zhang, Y. & Li, H. Signaling pathways and targeted therapies in lung squamous cell carcinoma: mechanisms and clinical trials. Signal Transduct Target Ther 7, 353, doi:10.1038/s41392-022-01200-x (2022).

29. Takahashi, K. & Yamanaka, S. Induction of pluripotent stem cells from mouse embryonic and adult fibroblast cultures by defined factors. Cell 126, 663–676, doi:10.1016/j.cell.2006.07.024 (2006).

30. Nakatsugawa, M. et al. SOX2 is overexpressed in stem-like cells of human lung adenocarcinoma and augments the tumorigenicity. Lab Invest 91, 1796–1804, doi:10.1038/labinvest.2011.140 (2011).

31. Singh, S., et al. EGFR/Src/Akt signaling modulates Sox2 expression and self-renewal of stem-like side-population cells in non-small cell lung cancer. Mol Cancer 11, 73, doi:10.1186/1476-4598-11-73 (2012).

32. Herbst, R. S., Morgensztern, D. & Boshoff, C. The biology and management of non-small cell lung cancer. Nature 553, 446–454, doi:10.1038/nature25183 (2018).

33. Gabasa, M., et al. MMP1 drives tumor progression in large cell carcinoma of the lung through fibroblast senescence. Cancer Lett 507, 1–12, doi:10.1016/j.canlet.2021.01.028 (2021).

34. Wang, B. Y. et al. The comparison between adenocarcinoma and squamous cell carcinoma in lung cancer patients. J Cancer Res Clin Oncol 146, 43–52, doi:10.1007/s00432-019-03079-8 (2020).

35. Saeed, A. M. et al. The influence of Hispanic ethnicity on nonsmall cell lung cancer histology and patient survival: an analysis of the Survival, Epidemiology, and End Results database. Cancer 118, 4495–4501, doi:10.1002/cncr.26686 (2012).

36. Friedlaender, A., Perol, M., Banna, G. L., Parikh, K. & Addeo, A. Oncogenic alterations in advanced NSCLC: a molecular super-highway. Biomark Res 12, 24, doi:10.1186/s40364-024-00566-0 (2024).

37. de Jager, V. D. et al. Future perspective for the application of predictive biomarker testing in advanced stage non-small cell lung cancer. Lancet Reg Health Eur 38, 100839, doi:10.1016/j.lanepe.2024.100839 (2024).

38. Arbour, K. C. et al. Effects of Co-occurring Genomic Alterations on Outcomes in Patients with. Clin Cancer Res 24, 334–340, doi:10.1158/1078-0432.CCR-17-1841 (2018).

39. Thummalapalli, R. et al. Clinical and Genomic Features of Response and Toxicity to Sotorasib in a Real-World Cohort of Patients With Advanced. JCO Precis Oncol 7, e2300030, doi:10.1200/PO.23.00030 (2023).

40. Binkley, M. S. et al. Mutations Predict Lung Cancer Radiation Resistance That Can Be Targeted by Glutaminase Inhibition. Cancer Discov 10, 1826–1841, doi:10.1158/2159-8290.CD-20-0282 (2020).

41. West, H. J. et al. Clinical efficacy of atezolizumab plus bevacizumab and chemotherapy in KRAS-mutated non-small cell lung cancer with STK11, KEAP1, or TP53 comutations: subgroup results from the phase III IMpower150 trial. J Immunother Cancer 10, e003027, doi:10.1136/jitc-2021-003027 (2022).

42. Choy, L. et al. Constitutive NOTCH3 Signaling Promotes the Growth of Basal Breast Cancers. Cancer Res 77, 1439–1452, doi:10.1158/0008-5472.CAN-16-1022 (2017).

43. Lin, S. et al. Non-canonical NOTCH3 signalling limits tumour angiogenesis. Nat Commun 8, 16074, doi:10.1038/ncomms16074 (2017).

44. Alam, M. M. et al. Contribution of NRF2 to sulfur metabolism and mitochondrial activity. Redox Biol 60, 102624, doi:10.1016/j.redox.2023.102624 (2023).

45. Honda, K. et al. On-tissue polysulfide visualization by surface-enhanced Raman spectroscopy benefits patients with ovarian cancer to predict post-operative chemosensitivity. Redox Biol 41, 101926, doi:10.1016/j.redox.2021.101926 (2021)

46. Erdélyi, K. et al. Reprogrammed transsulfuration promotes basal-like breast tumor progression via realigning cellular cysteine persulfidation. Proc Natl Acad Sci U S A 118, e2100050118, doi:10.1073/pnas.2100050118<otherinfo> (2021)</otherinfo>.

47. Takeda, H. et al. Sulfur metabolic response in macrophage limits excessive inflammatory response by creating a negative feedback loop. Redox Biol 65, 102834, doi:10.1016/j.redox.2023.102834 (2023).

48. Sekine, H. et al. PNPO-PLP axis senses prolonged hypoxia in macrophages by regulating lysosomal activity. Nat Metab 6, 1108–1127, doi:10.1038/s42255-024-01053-4 (2024).

49. Visvader, J. E. & Lindeman, G. J. Cancer stem cells: current status and evolving complexities. Cell Stem Cell 10, 717–728, doi:10.1016/j.stem.2012.05.007 (2012).

50. Cui, X., et al. Regnase-1 and Roquin Nonredundantly Regulate Th1 Differentiation Causing Cardiac Inflammation and Fibrosis. J Immunol 199, 4066–4077, doi:10.4049/jimmunol.1701211 (2017).

51. Yang, G., et al. Regnase-1 plays an essential role in maintaining skin immune homeostasis via regulation of chemokine expression. Biomed Pharmacother 162, 114558, doi:10.1016/j.biopha.2023.114558 (2023).

52. Htun, T. S., Tanaka, H., Singh, S. K., Diez, D. & Akira, S. Regnase-1 D141N mutation induces CD4+ T cell-mediated lung granuloma formation via upregulation of Pim2. Int Immunol 36, 497–516, doi:10.1093/intimm/dxae026 (2024).

53. Uehata, T. et al. Malt1-induced cleavage of regnase-1 in CD4(+) helper T cells regulates immune activation. Cell 153, 1036–1049, doi:10.1016/j.cell.2013.04.034 (2013).

54. Niu, J., Azfer, A., Zhelyabovska, O., Fatma, S. & Kolattukudy, P. E. Monocyte chemotactic protein (MCP)-1 promotes angiogenesis via a novel transcription factor, MCP-1-induced protein (MCPIP). J Biol Chem 283, 14542–14551, doi:10.1074/jbc.M802139200 (2008).

55. Caldwell, G. M., et al. The Wnt antagonist sFRP1 in colorectal tumorigenesis. Cancer Res 64, 883–888, doi: 10.1158/0008-5472.can-03-1346 (2004).

56. Sanjana, N. E., Shalem, O. & Zhang, F. Improved vectors and genome-wide libraries for CRISPR screening. Nat Methods 11, 783–784, doi:10.1038/nmeth.3047 (2014).

57. Yoshimachi, S. et al. Ral GTPase-activating protein regulates the malignancy of pancreatic ductal adenocarcinoma. Cancer Sci 112, 3064–3073, doi:10.1111/cas.14970 (2021).

