## Supplementary Figures, Tables, and Methods for "Regnase-1 Promotes Tumor-Initiating Activity in Non-Small Cell Lung Cancer"

### Regnase-1 promotes cancer stem-like phenotype in non-small cell lung cancer

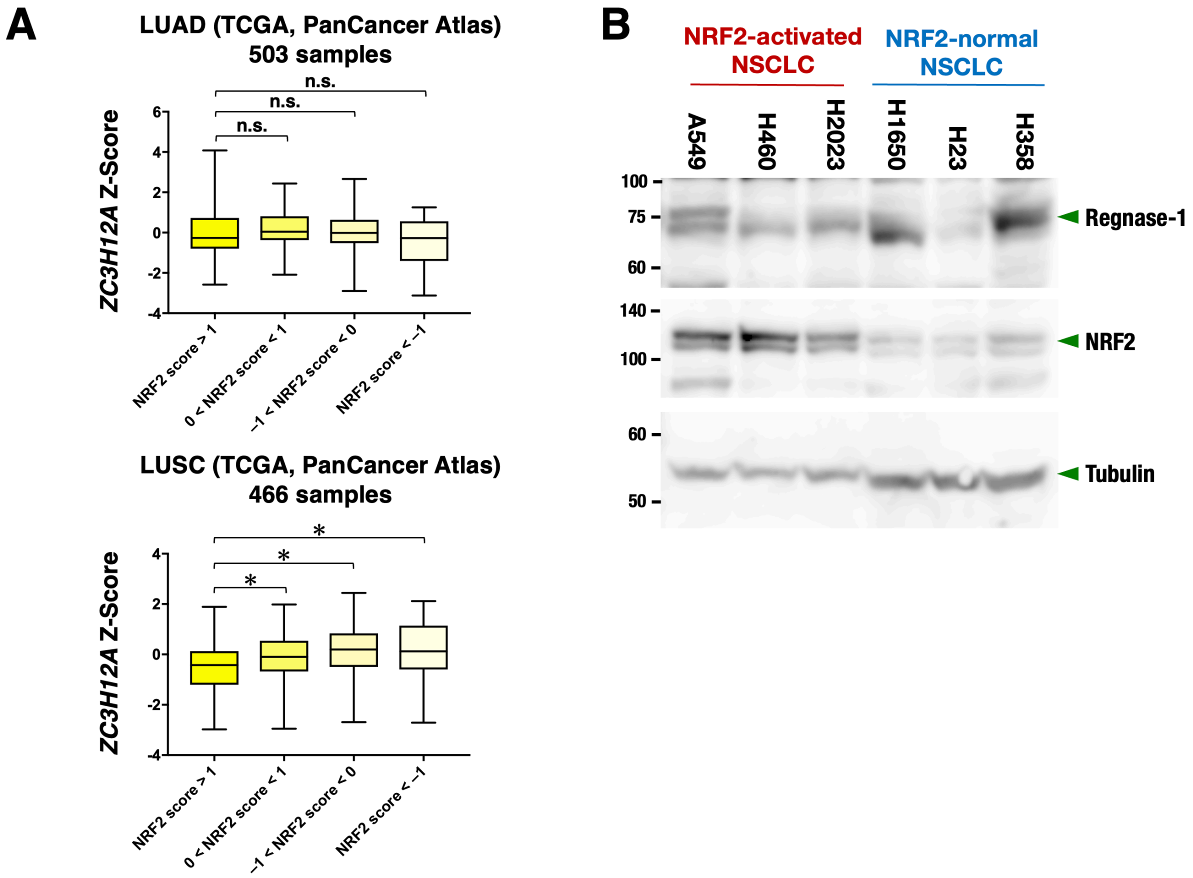


**Supplementary Figure S1.** ***ZC3H12A*** **expression is not associated with NRF2 activity.**

(**A**) *ZC3H12A* mRNA expression in tumor tissues stratified according to NRF2 activities. Gene expression data for patients with lung adenocarcinoma (LUAD) and lung squamous cell carcinoma (LUSC) from the TCGA database are shown. Box plots are defined by the 25th and 75th percentiles. Center line represents the median (50th percentile). Whiskers indicate minimum and maximum values. One-way ANOVA followed by the Bonferroni post hoc test was performed. *$p$<0.05, n.s.: not significant.

(**B**) Immunoblot analysis detecting Regnase-1 and NRF2 levels in NRF2-activated and NRF2-normal NSCLC cell lines. Tubulin was detected as a loading control. The result shown is a representative of 3 independent experiments.

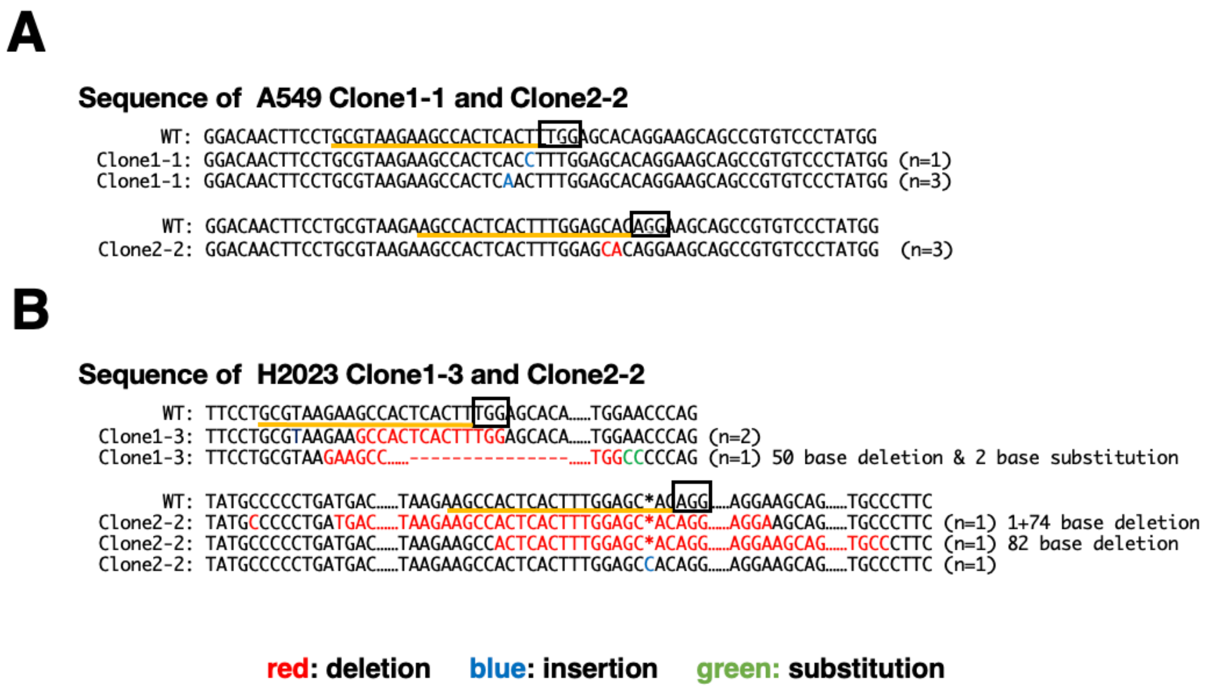


**Supplementary Figure S2. Genome editing of the *ZC3H12A* gene in A549 and H2023 cells.** DNA sequences were altered by CRISPR-Cas9 genome editing in the *ZC3H12A* locus of A549 (**A**) and H2023 (**B**) cells. Sequences corresponding to gRNAs are underlined in orange, and protospacer-adjacent motifs are boxed. PCR products were cloned and sequenced using primer pairs flanking the targeted regions. Deleted, inserted and substituted bases are indicated in red, blue and green, respectively. The numbers of altered sequences obtained in the cloned PCR products are shown to the right of each sequence.

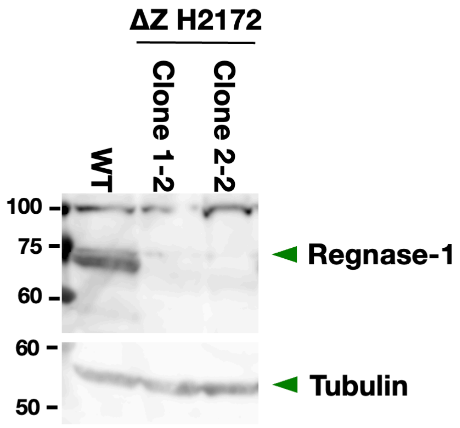


**Supplementary Figure S3. Establishment of ΔZ cells in H2172 cells.**

Immunoblot analysis detecting Regnase-1 in WT (wild-type) and *ZC3H12A*-deficient (ΔZ) H2172 cells. Tubulin was detected as a loading control. The result shown is a representative of 3 independent experiments.

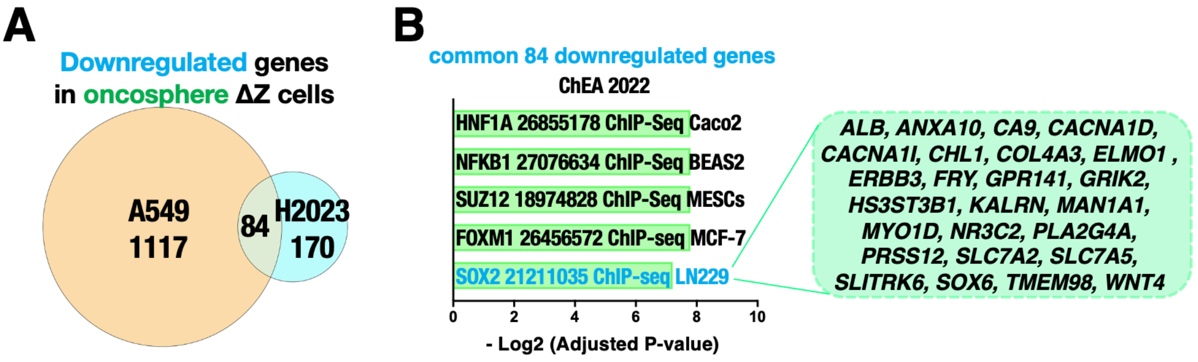


**Supplementary Figure S4. Regnase-1 dependent transcriptome in the oncosphere culture condition.**

(**A**) Regnase-1-dependent transcriptome obtained from RNA-seq analysis of A549 and H2023 cells in the oncosphere culture condition. Downregulated genes in ΔZ cells were defined as significantly downregulated genes (adjusted P value<0.05) in Clone 2-2 ΔZ vs. WT of A549 cells and Clone 2-2 ΔZ vs. WT of H2023 cells. 84 genes were identified as commonly downregulated genes between A549 and H2023 cells.

(**B**) Annotation analysis of the commonly downregulated genes was performed using ‘Enrichr’. The pathways shown in light blue are significantly decreased in ΔZ cells. Statistical significance was defined as an adjusted P-value of < 0.05.

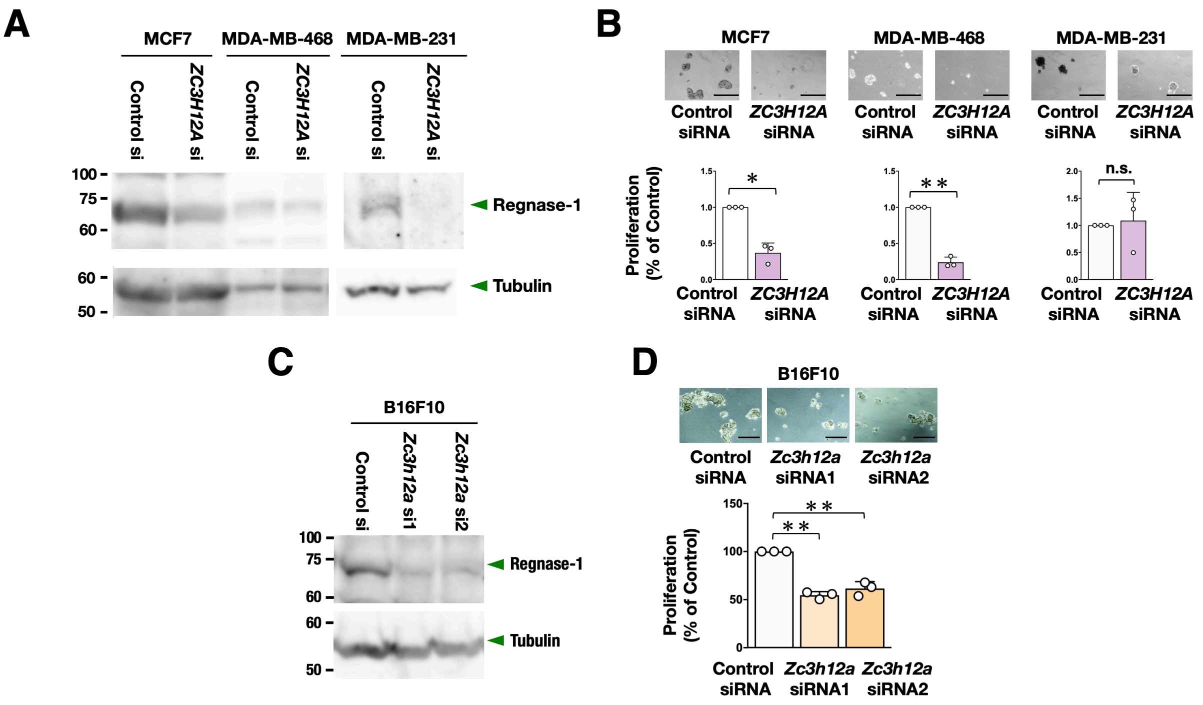


**Supplementary Figure S5. Regnase-1 regulates stem-like phenotype in cancer cell lines other than NSCLC.**

(**A, C**) Immunoblot analysis detecting Regnase-1 levels in breast cancer cell lines (**A**) and a murine melanoma cell line (**C**) treated with *ZC3H12A* siRNA (**A**) and *Zc3h12a* siRNAs (**C**) together with control siRNA, respectively. Tubulin was detected as a loading control.

(**B, D**)**.** Oncosphere growth of breast cancer cell lines treated with control siRNA and *ZC3H12A* siRNA (**B**) and a murine melanoma cell line treated with control siRNA and *Zc3h12a* siRNAs (**D**). Images of oncosphere growth treated with control siRNA and *ZC3H12A* siRNA or *Zc3h12a* siRNAs (upper panels) and viable cell counts after trypsinization (lower panels). Fold changes were calculated in comparison to control siRNA. Average cell numbers and SD from 3 independent experiments are shown. Two-sided confidence interval estimation was conducted to evaluate statistical significance. * $\alpha<0.05$, ** $\alpha<0.01$, n.s.: not significant. Scale bars indicate 200 μm for MDA-MB-231 and MDA-MB-468 cells, 500 μm for MCF7 and B16F10 cells.

**Supplementary Table S1. Oligo DNAs for construction of gRNA expression vectors to disrupt *ZC3H12A*.**

| Name | Sequence (5'-3') |
|---|---|
| ZC3H12A gRNA1 forward | CACCGGCGTAAGAAGCCACTCACTT |
| ZC3H12A gRNA1 reverse | AAACAAGTGAGTGGCTTCTTACGCC |
| ZC3H12A gRNA2 forward | CACCGAGCCACTCACTTTGGAGCAC |
| ZC3H12A gRNA2 reverse | AAACGTGCTCCAAAGTGAGTGGCTC |

**Supplementary Table S2. Primer set for the PCR Amplification of the DNA fragment spanning the gRNA target site.**

| Name | Sequence (5’-3’) |
|---|---|
| *ZC3H12A* PCR forward # | CTCCCACCTCCAGAGAGTTC |
| *ZC3H12A* PCR reverse # | AGAACCAGGGATGAGAGCAG |

#### Supplementary Methods Doc. S1

##### Cell culture.

NSCLC cell lines (A549, NCIH2023, NCIH2172, NCIH1944, NCIH1437, NCIH838, NCIH1650, NCIH23, NCIH358, CORL105, HCC4006, LK2, EBC1, SKMES1, NCIH460, LU65), breast cancer cell lines (MCF7, MDA-MB-468, DMA-MB-231), murine melanoma cell line (B16F10) were used in this study. All cell lines were maintained in low-glucose DMEM supplemented with 10% fetal bovine serum (FBS) and penicillin/streptomycin (Gibco). The cells were maintained in a 5% $CO_2$ atmosphere at 37 °C. PCR was used to confirm that the cultured cells were not infected with mycoplasma.

##### Immunoblot analyses.

For the preparation of whole-cell lysates, cells were directly lysed in 2x Laemmli buffer followed by boiling at 95 °C for 10 min. The protein samples were separated by SDS-PAGE and transferred onto PVDF membranes (Immobilon P, Millipore). The antibodies used were as follows: anti-NRF2 (sc-13032X, Santa Cruz), anti-Regnase-1 antibody (HPA032053, ATLAS Antibodies), and anti-tubulin (T9026, Sigma). The immunoblotting results were visualized with Chemi-Lumi One L (Nacalai Tesque) or ECL Prime (GE Healthcare) using Amersham ImageQuant 800 (Cytiva).

##### RNA-sequencing analysis.

Total RNA was extracted from WT/Clone2-2 A549 and WT/Clone2-2 H2023 cells using the RNeasy Mini Kit (Qiagen). Cells were analyzed in biological duplicate. After performing quality control, libraries were constructed from total RNA using TruSeq Stranded mRNA LT Sample Prep Kit according to the manufacturer's protocol (TruSeq Stranded mRNA Sample Preparation Guide, Part # 15031047 Rev. E). Briefly, the sequencing library was created by randomly fragmenting the cDNA sample, followed by 5′ and 3′ adapter ligation. Adapter-ligated fragments were then purified through PCR amplification and gel electrophoresis. The libraries were then sequenced on an Illumina platform, generating 102-base pair-end reads. The BCL (base calls) binary was then transformed into FASTQ format using the bcl2fastq package developed by Illumina. All of the above procedures, except for RNA purification, were conducted by Macrogen Japan (Kyoto, Japan). After obtaining raw fastq files, fastp (version 0.20.1) was conducted to assess the quality and trim possible adapters, poly-A tails, and low-sequence quality bases. Then, the remaining reads were aligned to the Grch38 reference genome using STAR version 2.7.8a. After mapping, RSEM version 1.3.3 was used to calculate raw read counts and the TPM (Transcripts Per Kilobase Million) value of each gene. After quantification, data were transferred to RStudio (ver. 2022.07.1 Build 554). Among the 60676 genes quantified in A549, those with extremely low counts (the Means of all samples were four or below) were eliminated, and 18361 genes were retained for further analysis. The same genes are selected for downstream analysis in H2023. A comparison between the two groups was made by the Wald-test using DEseq2 (ver. 1.34.0) with default parameters using raw read count data. For raw read count data, pseudo count (+1) was added to the raw read count of an individual gene. Genes with adjusted p values $< 0.05$ and $|\log_2 FC| > 1$ were considered DEGs. The Venn diagram using DEGs for each comparison was depicted using the "plotVenn" function. Enrichment analysis was performed on Enrichr (https://maayanlab.cloud/Enrichr/). The results were visualized as bar plots using ggplot2.

**Analysis of non-small cell lung cancer patients.**

***Correlation between ZC3H12A expression levels and prognosis.***

RNA-seq data of 498 lung adenocarcinoma (LUAD) and 493 lung squamous cell carcinoma (LUSC) cases from TCGA were analyzed. Overall survival was compared between two groups according to the FPKM value of *ZC3H12A*: the *ZCH3H12A* high-expressing group and the *ZCH3H12A* low-expressing group. The cut-off threshold was set at FPKM value 11.61 in LUAD, which can divide 140 samples for *ZC3H12A* high and 360 samples for *ZC3H12A* low. Also, the cut-off threshold was set at FPKM value 20.32 in LUSC, which can divide 108 samples for *ZC3H12A* high and 386 samples for *ZC3H12A* low.

***Meta-analysis of the TCGA database.***

RNA-seq data of 503 LUAD and 466 LUSC cases from TCGA, PanCancer Atlas were analyzed. NRF2 scores were defined as average values of z-scores of 6 representative NRF2 target genes, *NQO1*, *SLC7A11*, *GCLC*, *GCLM*, *TXNRD1* and *NR0B1*.[8]

**Transient knockdown experiments.**

*ZC3H12A* pooled siRNA against human Regnase-1 was purchased from Dharmacon (siGENOME SMARTpool siRNA M-014576-01-0010). *Zc3h12a* siRNAs against mouse Regnase-1 was purchased from Sigma (SASI_Mm01_0015-4747 and SASI_Mm01_0015-4748). MISSION siRNA Universal Negative Controls (Sigma-Aldrich) were used as controls. siRNAs were transfected into cells using Lipofectamine™ RNAiMAX Transfection Reagent (Thermo Fisher Scientific). Culture media were changed 24 hrs after transfection. After another 24-48 hrs, the cells were harvested for immunoblot analysis. The protocols used for transient knockdown with oncosphere formation assays in each cell line are described below.

**Establishment of inducible *ZC3H12A* knockdown cells.**

An inducible *ZC3H12A* shRNA lentiviral vector (Individual SMART vector Human Inducible Lentiviral shRNA; V3SH11252_226692445) and a control vector (SMART vector Inducible Non-targeting mCMV-TurboGFP; VSC11651) were purchased from Dharmacon. H460 cells and LK2 cells were infected with lentiviral particles with 12.5 μg/ml polybrene. After 24 hrs, the cells were replated in 10-cm dishes and cultured in selection medium containing 2 μg/ml puromycin. Single clones were selected using cloning rings (TOHO). Doxycycline (DOX) (1 μg/ml) or the same dose of DMSO was added 48 hrs prior to cell harvest for immunoblot analysis.

**Spheroid formation assay.**

Cell growth was examined in spheroid culture. $10^3$ cells of each cell line with 100 μl culture media were seeded in a low-attachment U-bottom plate (PrimeSurface 96 well Plate, MS-9096UZ, Sumitomo Bakelite Co.). Spheroids were observed 96 hrs after transfection. After 96 hrs, cell proliferation was assessed using the Cell Counting Kit-8 (Nacalai Tesque) according to the manufacturer's protocol.

**Oncosphere formation assay.**

Oncosphere formation was examined as one of the indicators of tumor-initiating activity. Briefly, cells were cultured in ultralow attachment dishes (Corning) in CSC medium. The CSC medium consisted of serum-free DMEM-F12 medium (Gibco-Invitrogen) containing 50 μg/ml insulin (Sigma-Aldrich), 0.4% albumin bovine fraction Ⅴ (Sigma-Aldrich), N-2 Plus Media Supplement (R&D Systems), Gibco B-27 Supplement (Thermo Fisher Scientific), 20 ng/ml EGF (Pepro Tech) and 10 ng/ml bFGF (Pepro Tech). To compare oncosphere formation of WT and ΔZ cells, A549 and H2023 ($2 \times 10^4$ cells) and H2172 ($4 \times 10^4$ cells) were seeded in CSC medium. The culture media were left unchanged until the cell harvest on day 7 after seeding. Viable cells were counted by trypan blue staining. For introduction of siRNAs against *ZC3H12A* or *Zc3h12a*, transfection of the siRNA was performed under regular culture conditions. After 24 hrs, HCC4006 ($10 \times 10^4$ cells), MDA-MB-231 ($8 \times 10^4$ cells), H1944,

H1437, H838, CORL105, H358, H23, SKMES1, MCF7 and MDA-MB-468 (6 x $10^4$ cells), A549, H2023, H2172, H1650, EBC1 and B16F10 (4 x $10^4$ cells), LK2 and H460 (2 x $10^4$ cells) were reseeded in CSC medium. The culture media were left unchanged until the cell harvest on day 5 after reseeding. Viable cells were counted by trypan blue staining. For inducible *ZC3H12A* knockdown of H460 and LK2 cells, procedures were described in schemes of Figures 6B and 6C. The culture media were left unchanged until cell harvest, and 1 μg/ml DOX or the same dose of DMSO was added to the media. Viable cells were counted using trypan blue staining.

**Xenograft experiments.**

Cell suspensions were mixed with Matrigel (Corning) and subcutaneously injected into the flanks of four-week-old male Balb/c nu/nu mice. To compare tumorigenesis between WT and ΔZ cells, $1 \times 10^6$ (A549), $0.5 \times 10^6$ (H2023), and $0.5 \times 10^6$ (H2172) cells were injected for each group, and the resulting tumors were dissected and weighed after 38 days (A549), 35 days (H2023), and 46 days (H2172), respectively. For inducible *ZC3H12A* knockdown in H460 cells shown in Figure 6D, $1 \times 10^4$ cells pretreated with DOX (1 μg/ml) for 48 hours were subcutaneously injected. The recipient Balb/c nu/nu mice were continuously treated with 1 mg/ml DOX in drinking water containing 5% sucrose immediately after transplantation, and tumors were analyzed on day 26 post-transplantation. In another experiment involving inducible *ZC3H12A* knockdown in H460 and LK2 cells shown in Figure 6E, $1 \times 10^4$ cells without DOX pretreatment were subcutaneously injected. The recipient Balb/c nu/nu mice were treated with or without 1 mg/ml DOX in drinking water containing 5% sucrose after tumor formation, as indicated, and the resulting tumors were analyzed at the specified time points post-transplantation.

**Serial transplantation experiments.**

Tumors from the primary xenograft experiment were dissected from the primary recipient mice and chopped into small pieces under sterile conditions and incubated at 37°C for 1 hr in 5 ml low glucose DMEM supplemented with 10% fetal bovine serum (FBS) and penicillin/streptomycin containing 1 mg/mL collagenase type IV (Sigma-Aldrich, #C5138) and 100 µg/ml DNase I (SIGMA). The samples were incubated with 5 ml RBC lysis buffer (155 mM $NH_4Cl$, 15 mM $NaHCO_3$, 0.1 M EDTA, pH7.3) for 10 min, followed by filtration to remove debris. The samples were then incubated with anti-biotinylated CD31 (#13-0311-82, eBioscience) and CD45 (#13-0451-85, eBioscience) antibodies, followed by reaction with Dynabeads M-280 streptavidin (Thermo Fisher Scientific) to remove murine cells of endothelial and hematopoietic origins. To further remove the remaining murine cells, the samples were reacted with an FITC-conjugated anti-mouse MCH class I antibody (ab95572, Abcam), and human tumor cells were sorted as FITC-negative fraction by flow cytometry (BD FACSAria II, Becton Dickinson). Then, the sorted $5 \times 10^3$ WT and ΔZ A549 cells were transplanted into secondary recipient mice. The time course of the experiment is shown in Figure 4D.

**Quantification and statistical analysis.**

Statistical significance was evaluated using the Wilcoxon rank-sum test and one-way ANOVA followed by the Bonferroni post hoc test. Confidence intervals were calculated for all fold change evaluations. These analyses were performed using Microsoft Office Excel (Microsoft), Prism 7 (GraphPad Software, Inc.), and JMP Pro 13. $P<0.05$ and $\alpha<0.05$ (confidence interval) were considered statistically significant.
