## Supplementary material for "Regnase-1 Promotes Tumor-Initiating Activity in Non-Small Cell Lung Cancer": Uncropped blots

### Figure 2A

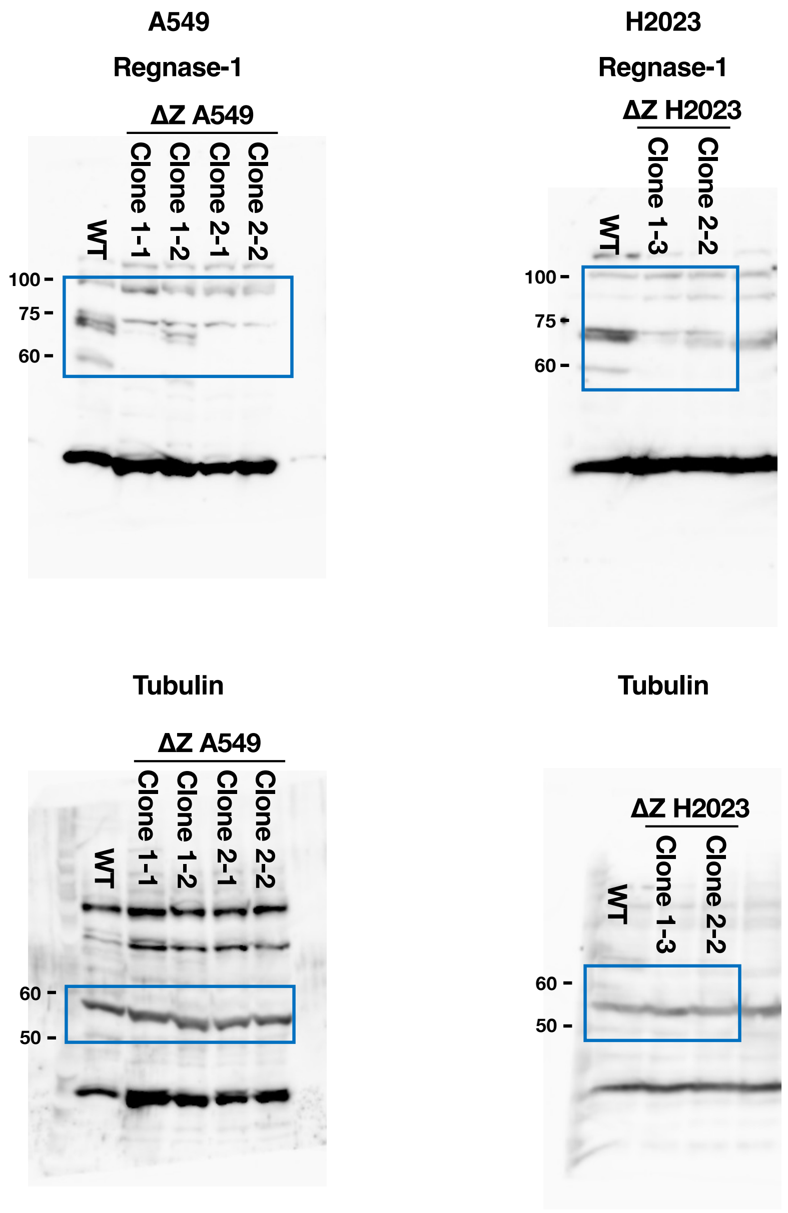


Uncropped blots shown in Figure 2A.
Areas surrounded by blue squares are regions of interest.

### Figure 2A
**Replicates data**

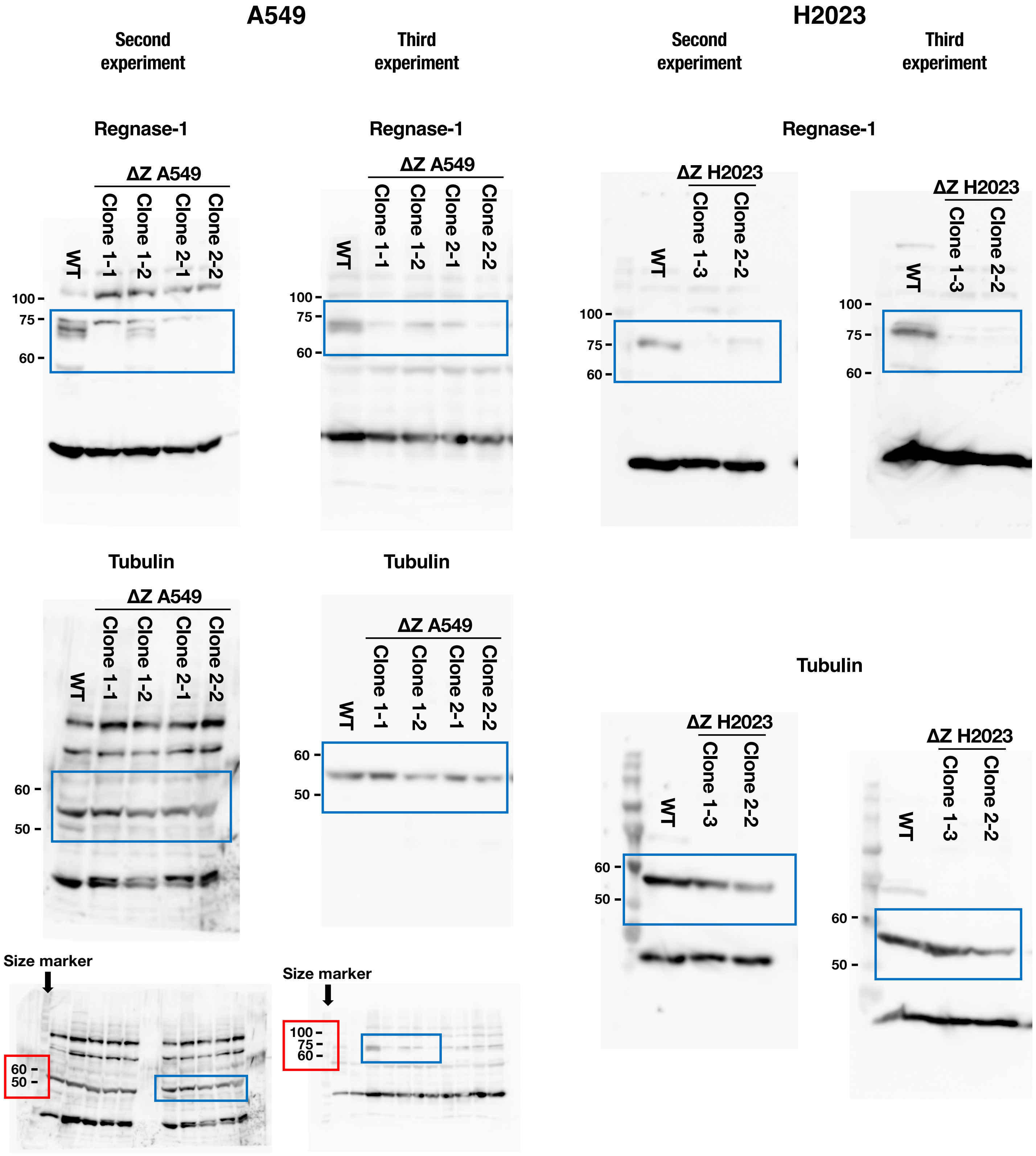


Uncropped blots of replicated data for those shown in Figure 2A. Areas surrounded by blue squares are regions of interest.
In cases where there are no size marker bands to the left of Regnase-1 or Tubulin, the raw data with the size markers are shown below. Areas surrounded by red squares are regions of interest size markers.

### Figure 5A

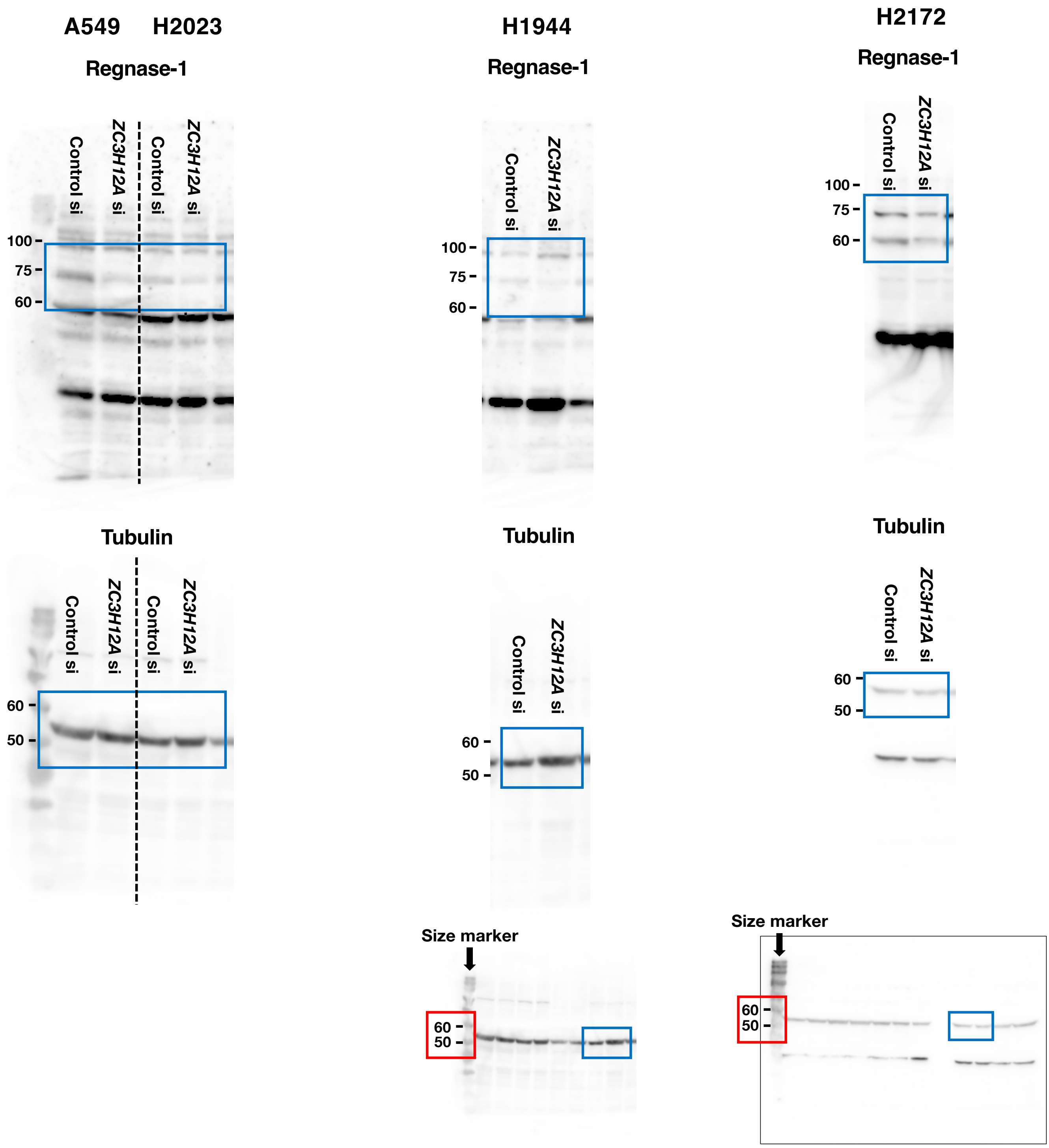


Uncropped blots of replicated data for those shown in Figure 5A. Areas surrounded by blue squares are regions of interest.
In cases where there are no size marker bands to the left of Regnase-1 or Tubulin, the raw data with the size markers are shown below. Areas surrounded by red squares are regions of interest size markers.
When the top and bottom ends of the membrane is unclear, the full length of the membrane is shown below and surrounded by a black line.

### Figure 5A

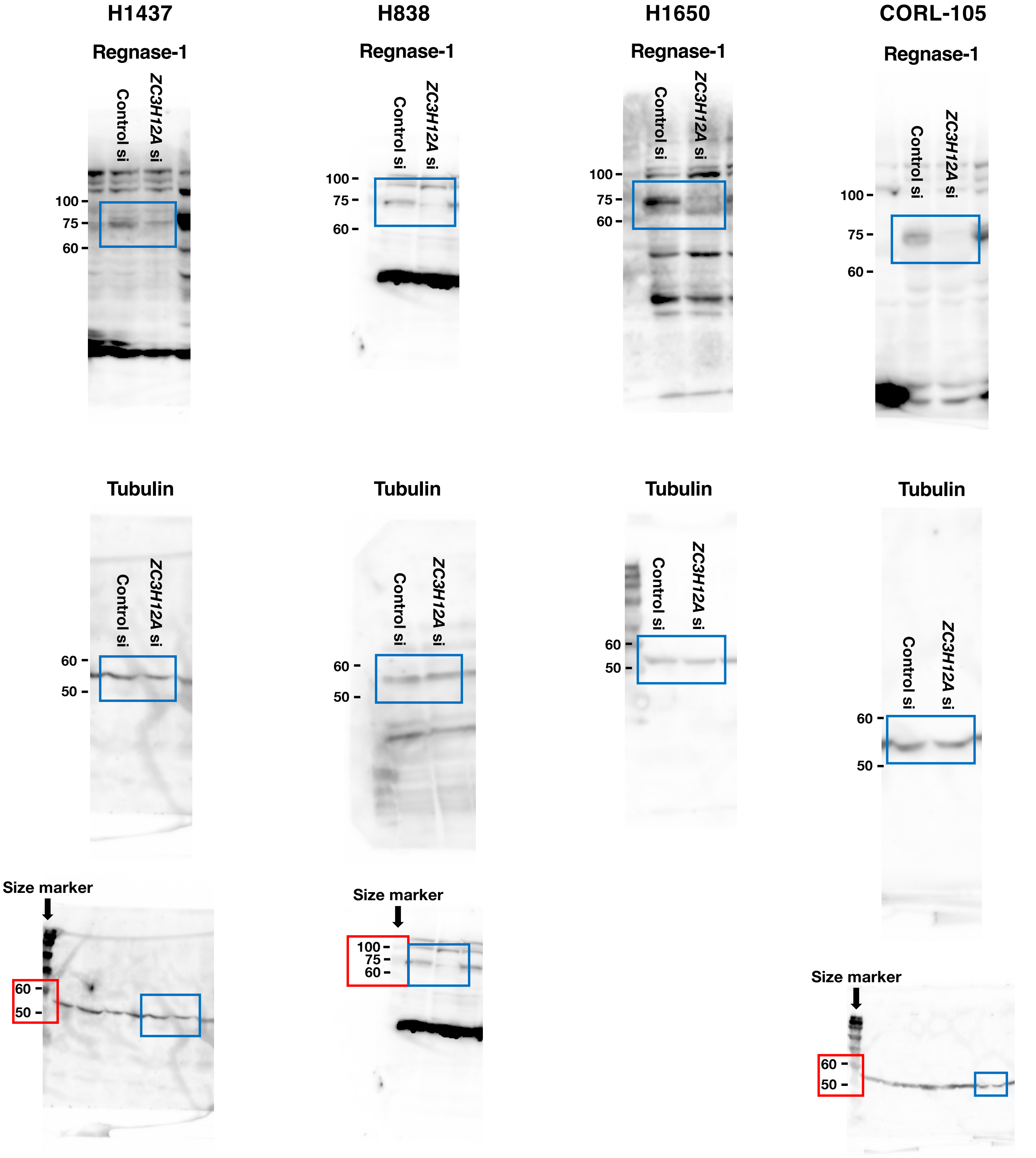


Uncropped blots of replicated data for those shown in Figure 5A. Areas surrounded by blue squares are regions of interest.
In cases where there are no size marker bands to the left of Regnase-1 or Tubulin, the raw data with the size markers are shown below. Areas surrounded by red squares are regions of interest size markers.

### Figure 5A

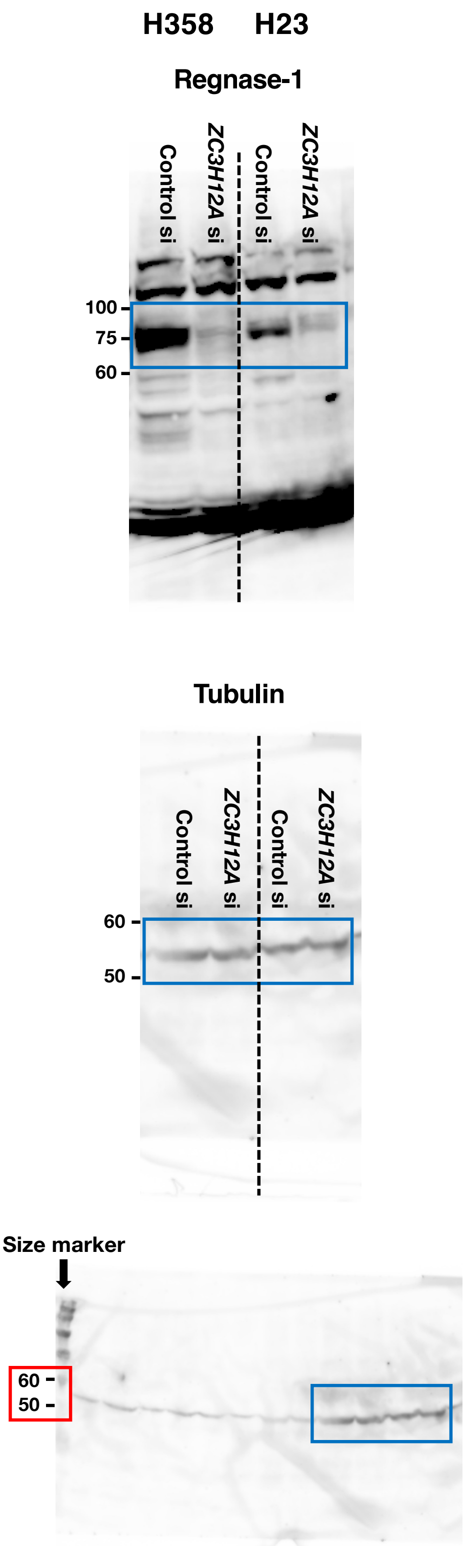


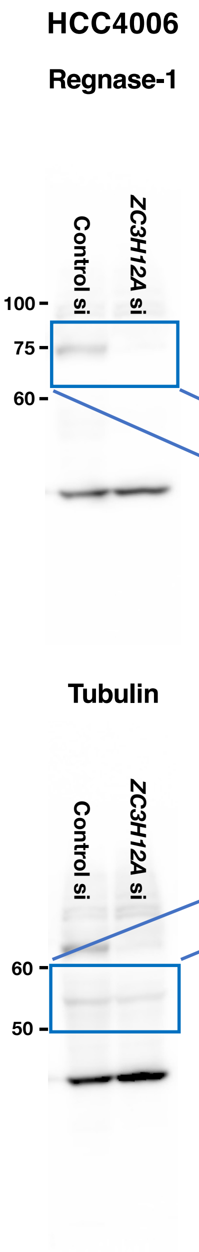


・Regnase-1: The same membrane was used with shorter exposure time than the whole blot below.

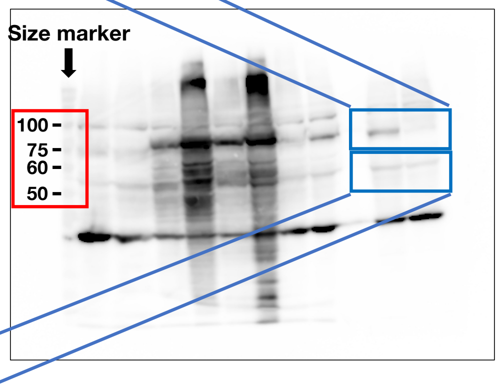


Uncropped blots of replicated data for those shown in Figure 5A. Areas surrounded by blue squares are regions of interest.
In case where there are no size marker bands to the left of Regnase-1 or Tubulin, the raw data with the size markers are shown below. Areas surrounded by red squares are regions of interest size markers.
When the top and bottom ends of the membrane are unclear, the full length of the membrane is shown right and surrounded by a black line.

### Figure 5A

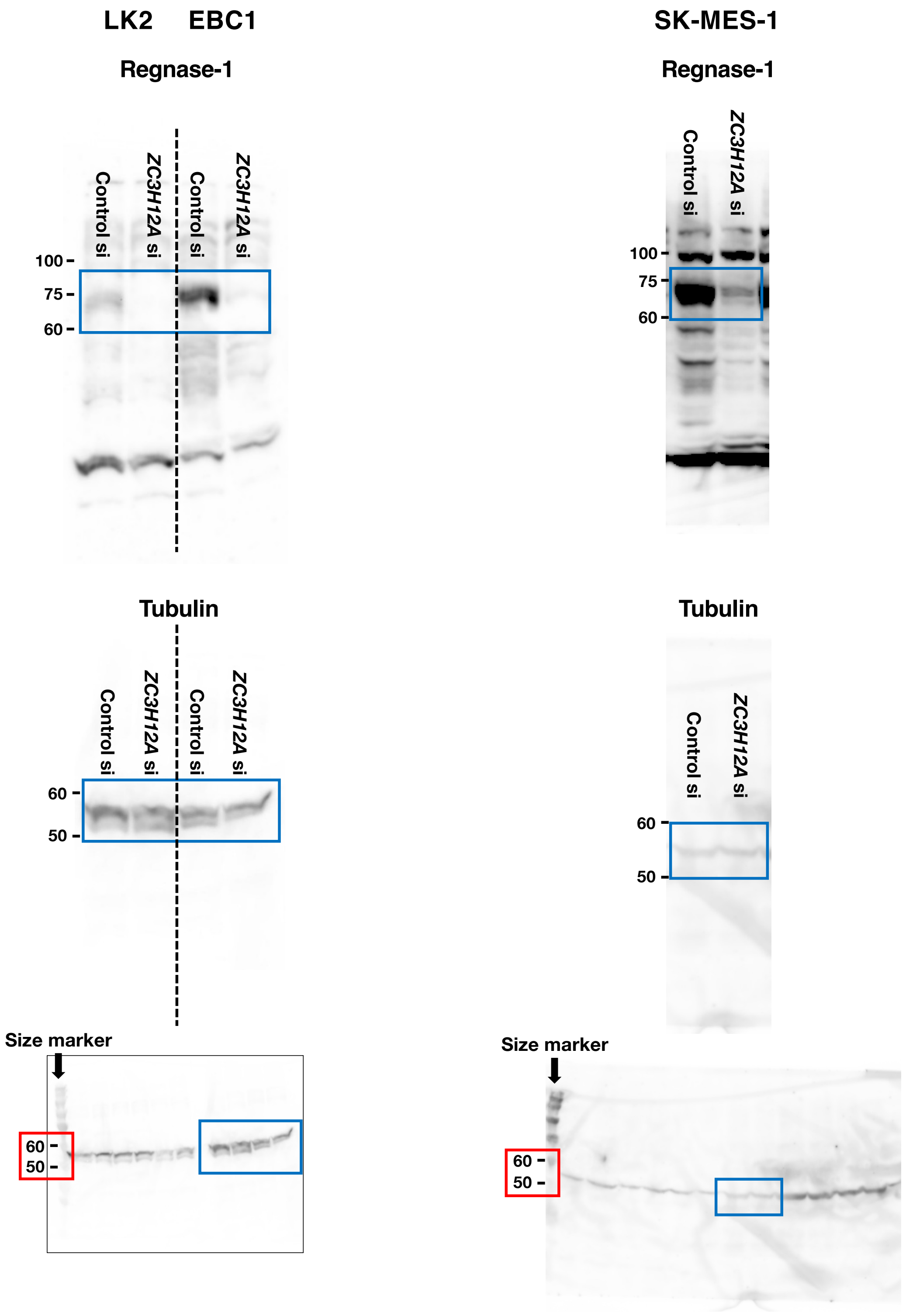


Uncropped blots of replicated data for those shown in Figure 5A. Areas surrounded by blue squares are regions of interest.
In case where there are no size marker bands to the left of Regnase-1 or Tubulin, the raw data with the size markers are shown below. Areas surrounded by red squares are regions of interest size markers.
When the top and bottom ends of the membrane are unclear, the full length of the membrane is shown below and surrounded by a black line.

### Figure 5A

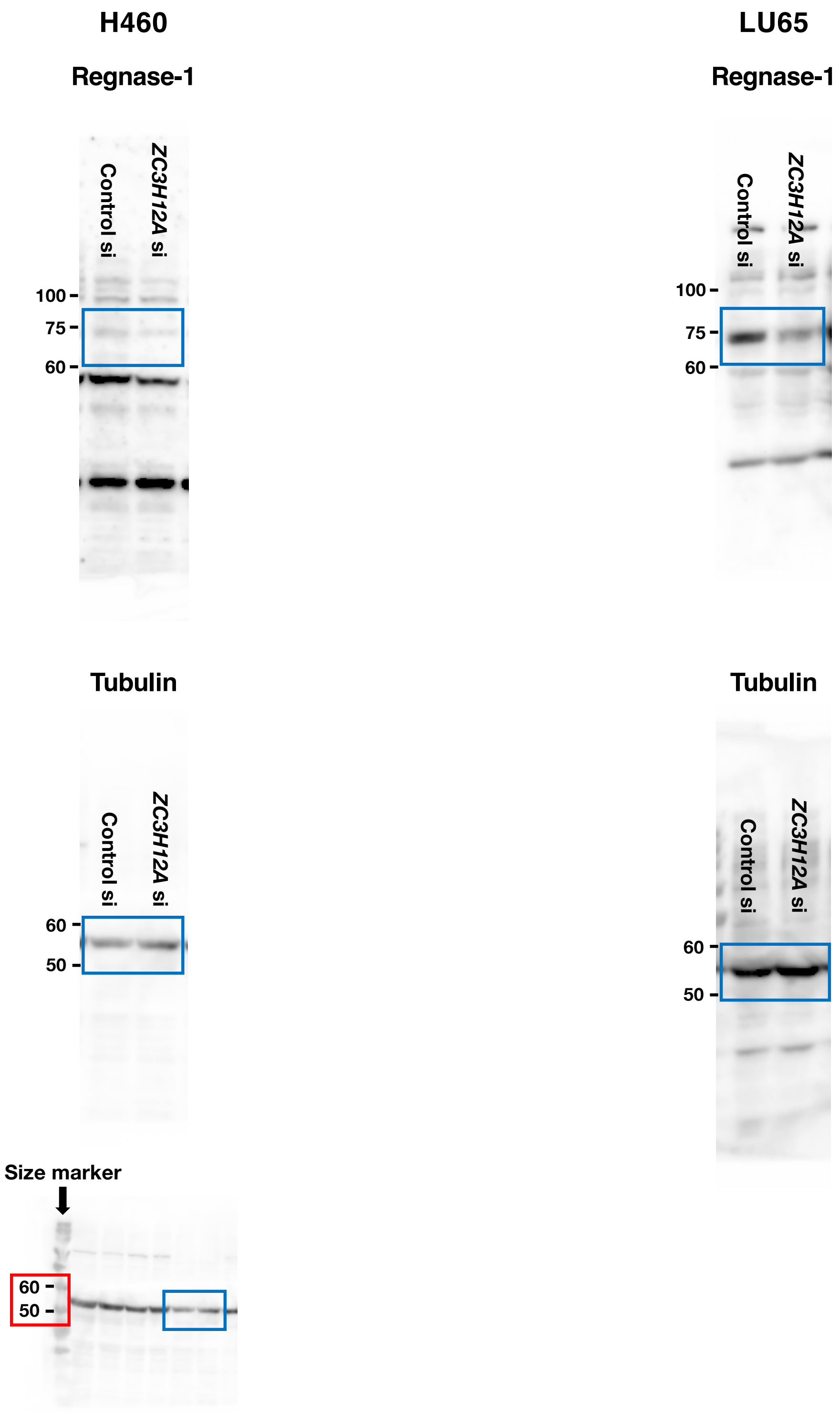


Uncropped blots of replicated data for those shown in Figure 5A. Areas surrounded by blue squares are regions of interest.
In cases where there are no size marker bands to the left of Regnase-1 or Tubulin, the raw data with the size markers are shown below. Area surrounded by red square is region of interest size markers.

### Figure 5A

**Replicate data for A549, EBC1, H460 and LU65**

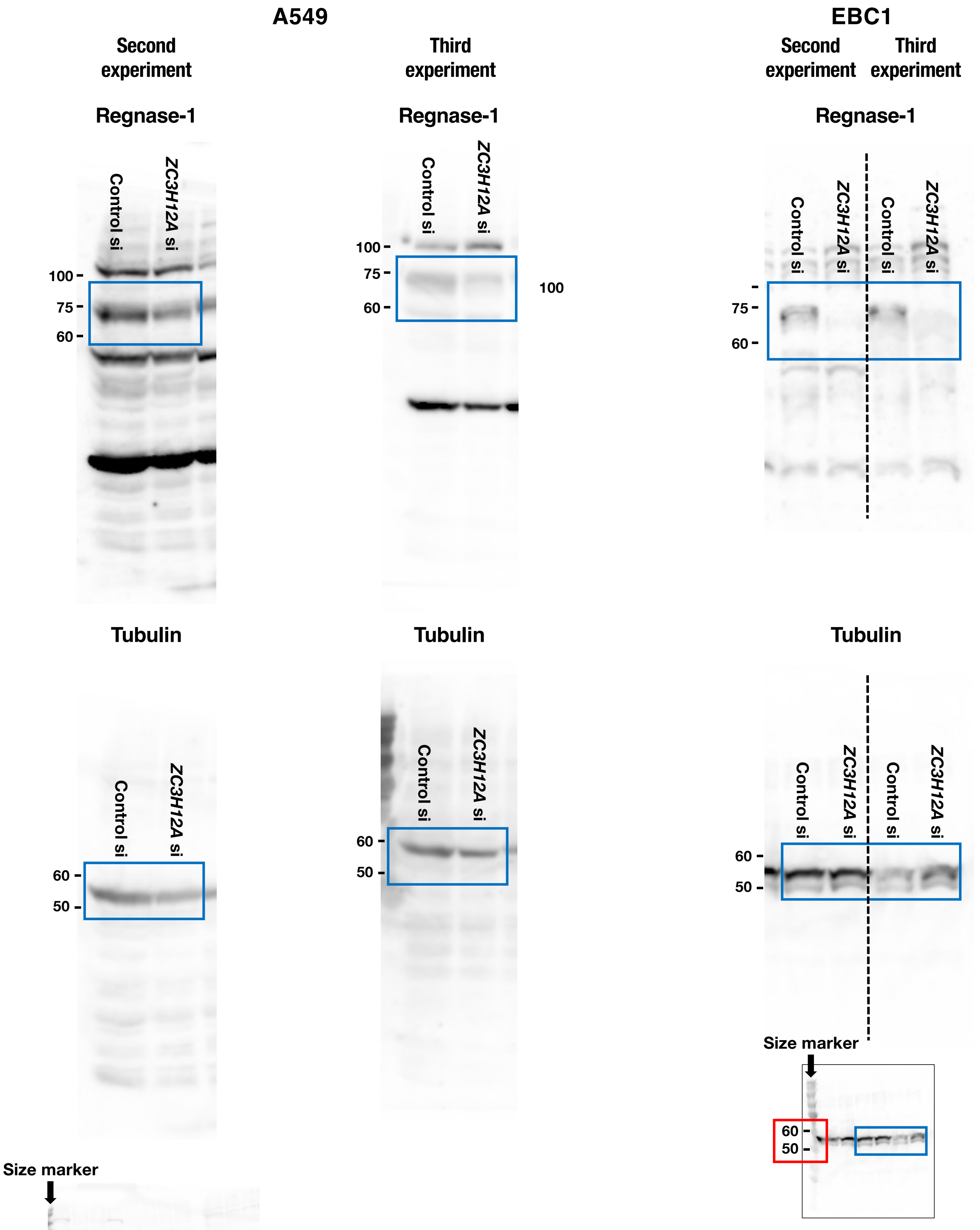


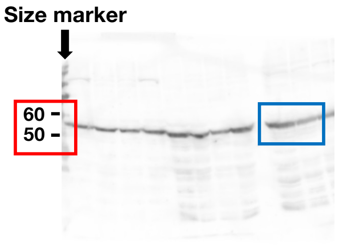


Uncropped blots of replicated data for those shown in Figure 5A. Areas surrounded by blue squares are regions of interest.
In cases where there are no size marker bands to the left of Regnase-1 or Tubulin, the raw data with the size markers are shown below. Areas surrounded by red squares are regions of interest size markers.
When the top and bottom ends of the membrane are unclear, the full length of the membrane is shown below and surrounded by a black line.

### Figure 5A

**Replicate data for A549, EBC1, H460 and LU65**

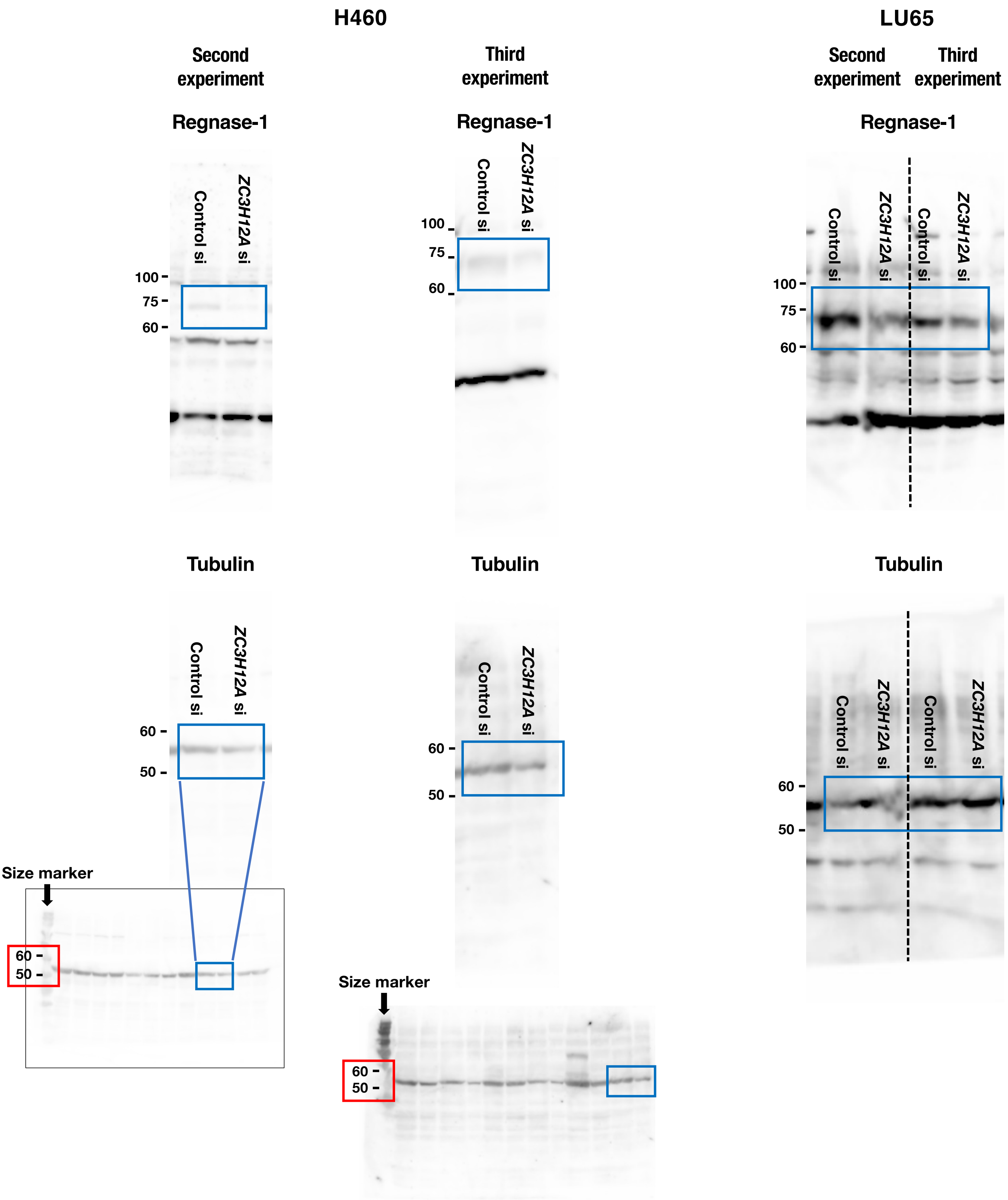


Uncropped blots of replicated data for those shown in Figure 5A. Areas surrounded by blue squares are regions of interest.

In cases where there are no size marker bands to the left of Regnase-1 or Tubulin, the raw data with the size markers are shown below. Areas surrounded by red squares are regions of interest size markers.

When the top and bottom ends of the membrane are unclear, the full length of the membrane is shown below and surrounded by a black line.

#### Figure 6A

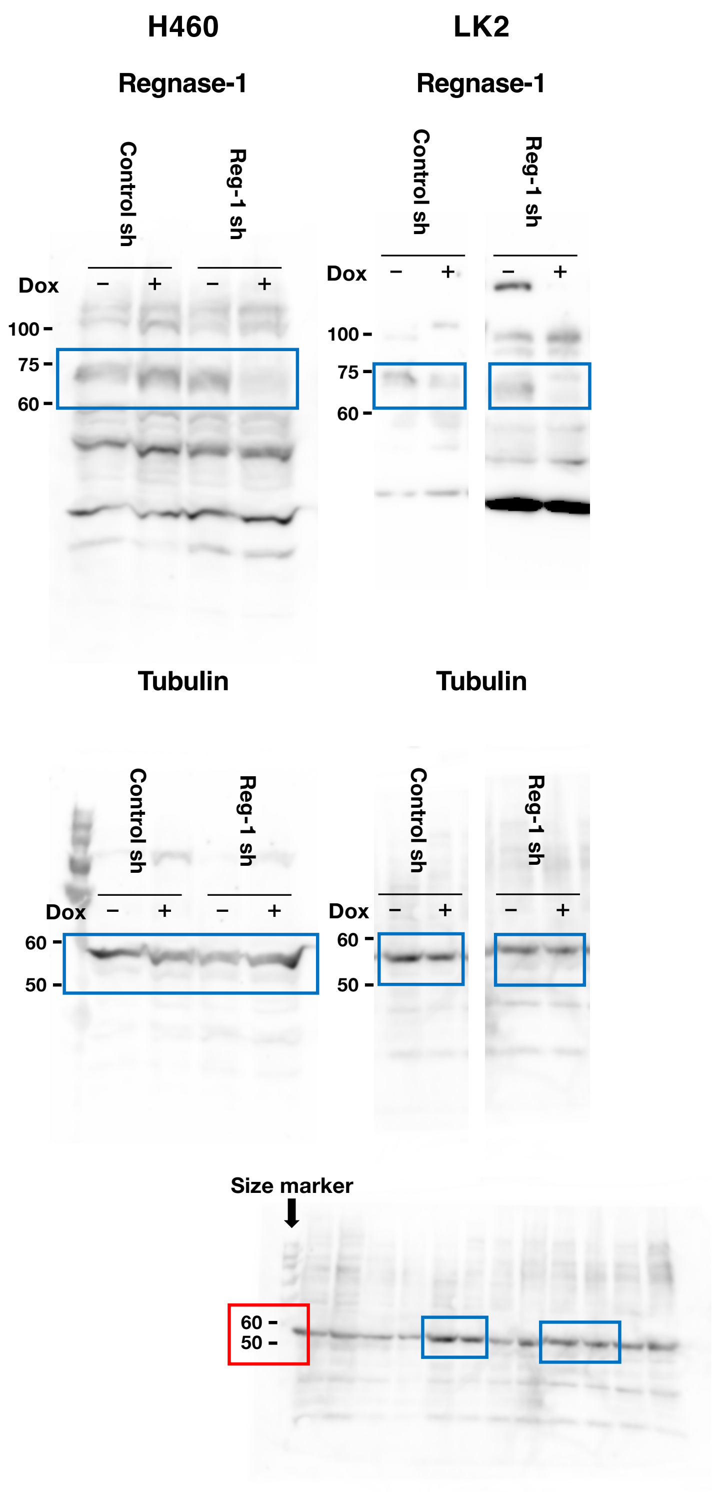


#### Figure 6A
**replicate data for H460 Reg-1 sh**

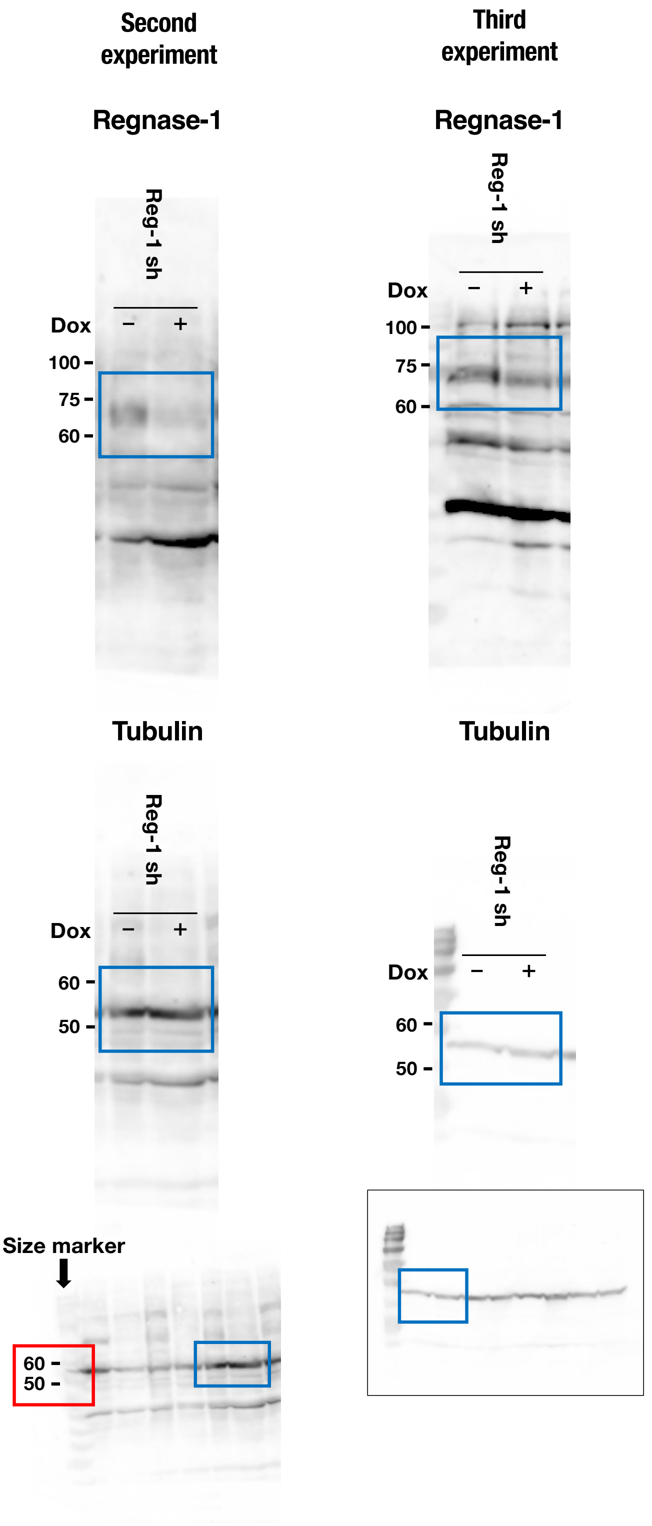


Uncropped blots of replicated data for those shown in Figure 6A. Areas surrounded by blue squares are regions of interest.
In cases where there are no size marker bands to the left of Regnase-1 or Tubulin, the raw data with the size markers are shown below. Areas surrounded by red squares are regions of interest size markers.
When the top and bottom ends of the membrane are unclear, the full length of the membrane is shown below and surrounded by a black line.

### Supplementary Figure S1B

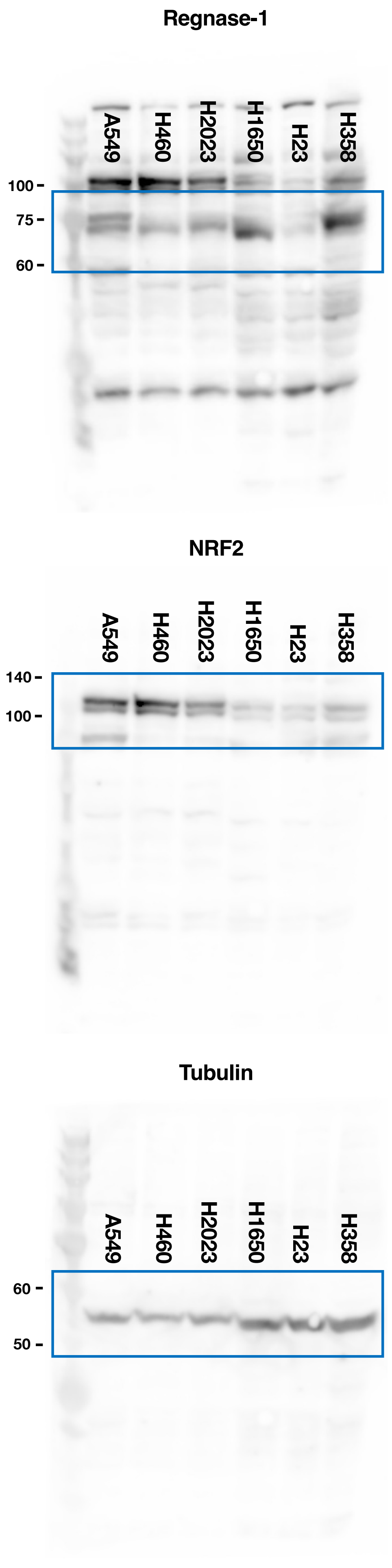


### Supplementary Figure S1B

#### Replicate data

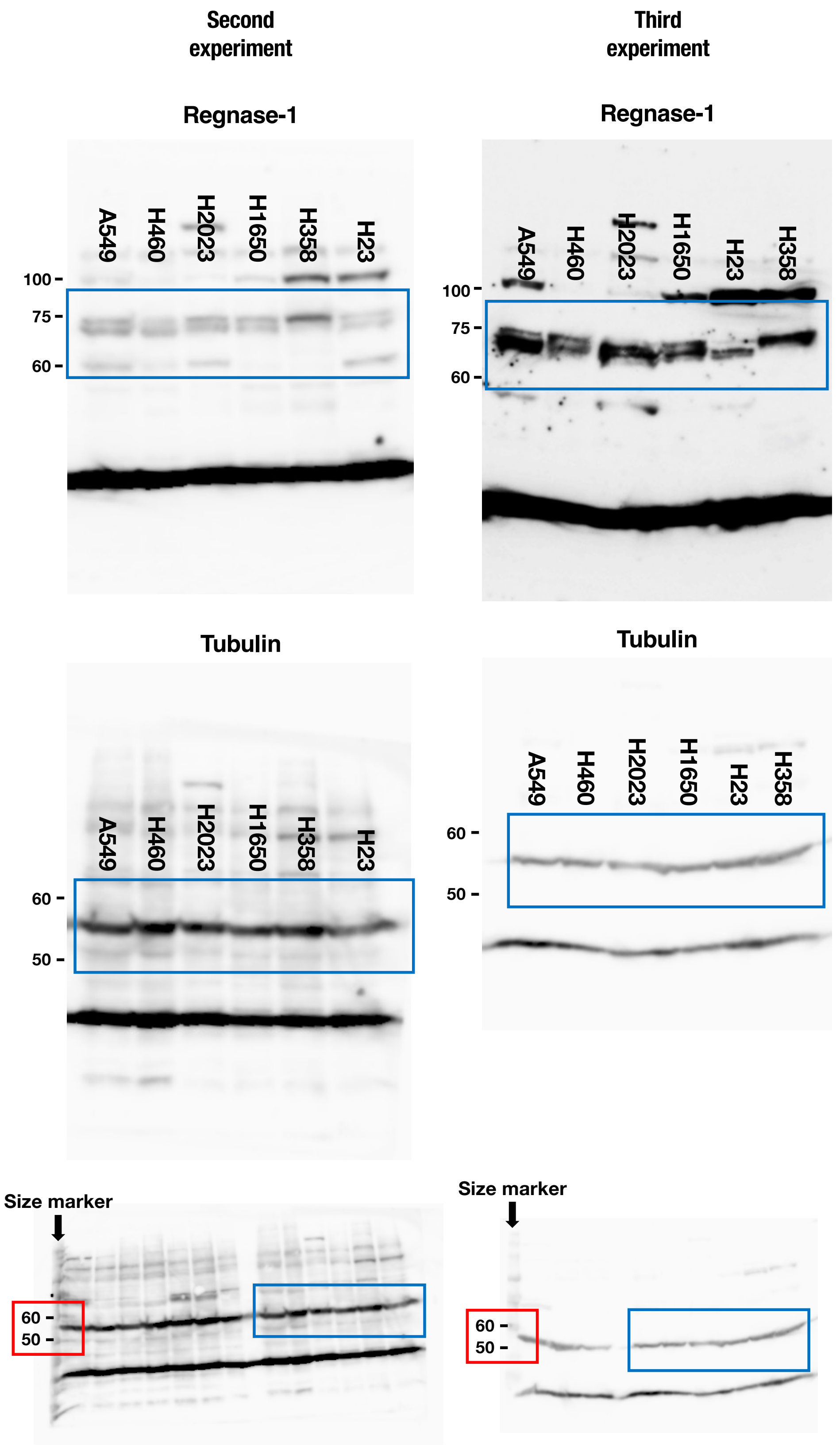


Uncropped blots of replicated data for those shown in Supplementary Figure S1B. Areas surrounded by blue squares are regions of interest.
In cases where there are no size marker bands to the left of Regnase-1 or Tubulin, the raw data with the size markers are shown below. Areas surrounded by red squares are regions of interest size markers.

#### Supplementary Figure S3

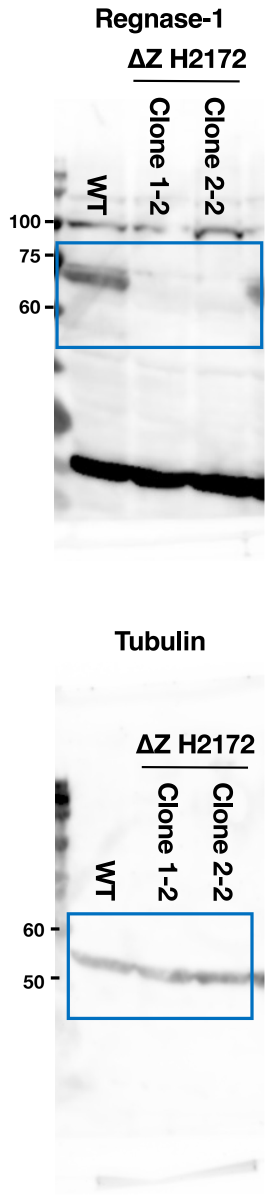


#### Supplementary Figure S3
**Replicates data**

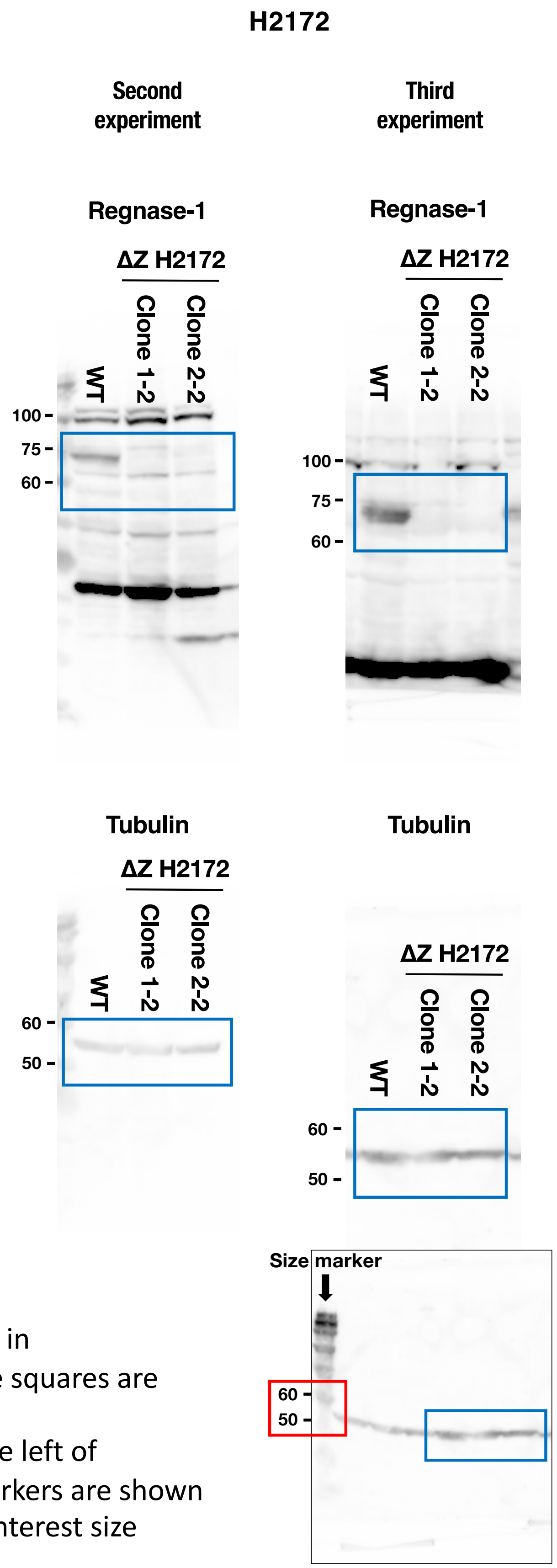


Uncropped blots of replicated data for those shown in Supplementary Figure S3. Areas surrounded by blue squares are regions of interest.
In cases where there are no size marker bands to the left of Regnase-1 or Tubulin, the raw data with the size markers are shown below. Area surrounded by red square is region of interest size markers.
When the top and bottom ends of the membrane are unclear, the full length of the membrane is shown below and surrounded by a black line.

### Supplementary Figure S5A

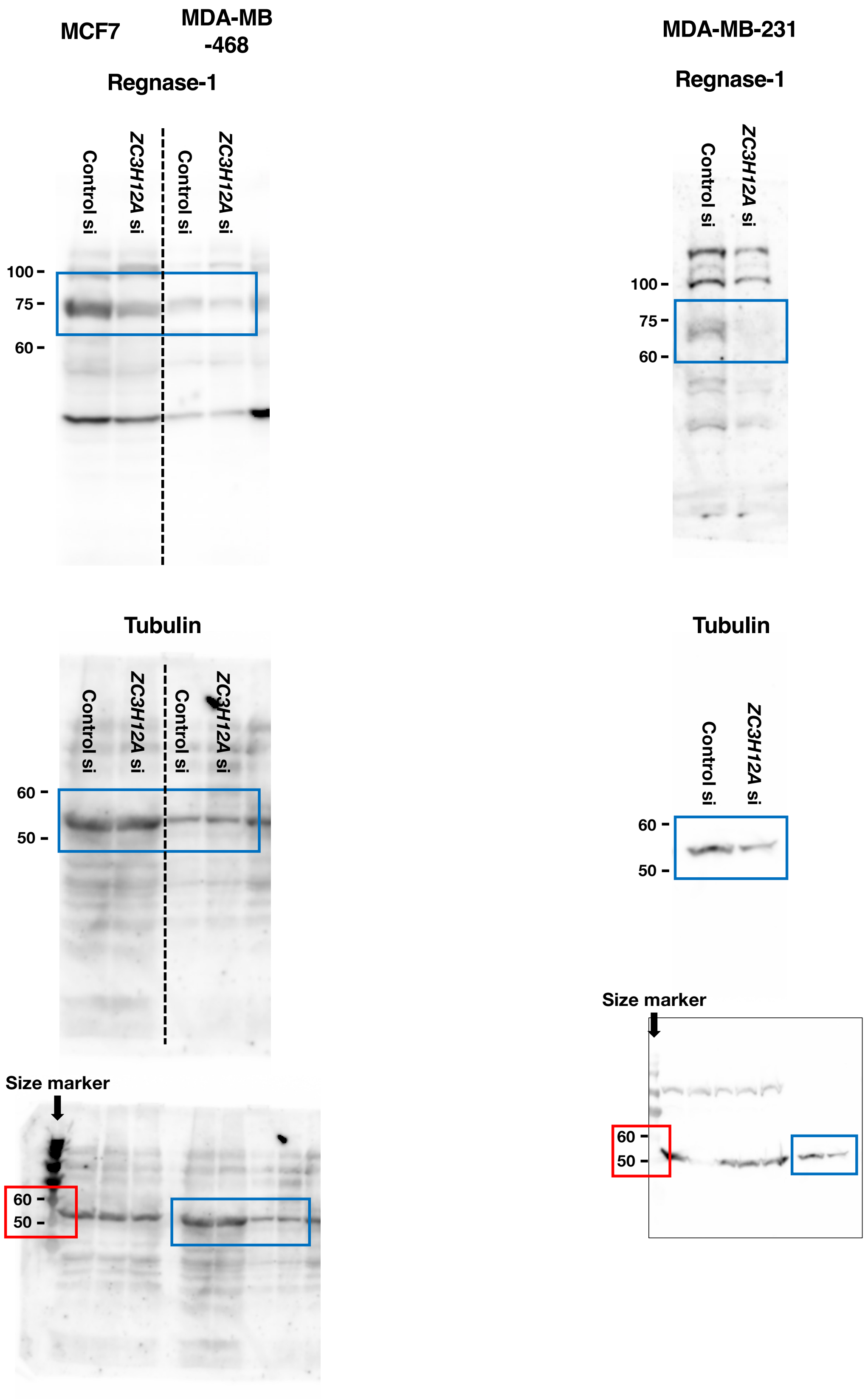


Uncropped blots of replicated data for those shown in Supplementary Figure S5A. Areas surrounded by blue squares are regions of interest.

In cases where there are no size marker bands to the left of Regnase-1 or Tubulin, the raw data with the size markers are shown below. Areas surrounded by red squares are regions of interest size markers.

When the top and bottom ends of the membrane are unclear, the full length of the membrane is shown below and surrounded by a black line.

#### Supplementary Figure S5C

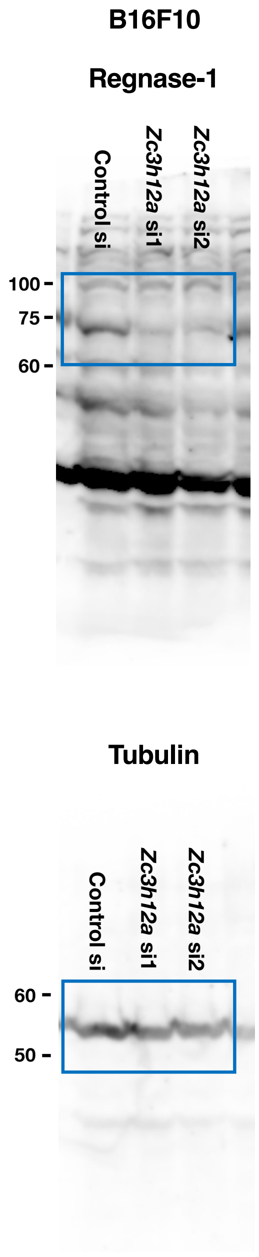


#### Supplementary Figure S5C

##### Replicates data

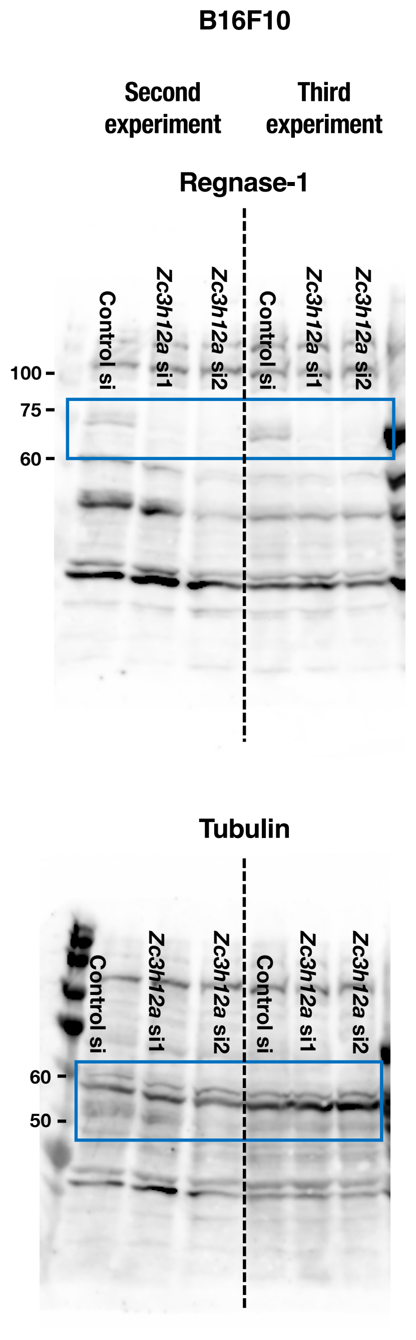


Uncropped blots shown in Supplementary Figure S5C and their replicated data. Areas surrounded by blue squares are regions of interest.
